# The evolution of reduced facilitation in a four-species bacterial community

**DOI:** 10.1101/2024.02.22.581583

**Authors:** Philippe Piccardi, Eric Ulrich, Marc Garcia-Garcerà, Rita Di Martino, Samuele E. A. Testa, Sara Mitri

## Abstract

Microbial evolution is typically studied in mono-cultures or in communities of competing species. But microbes do not always compete and how positive inter-species interactions drive evolution is less clear: Initially facilitative communities may either evolve increased mutualism, increased reliance on certain species according to the Black Queen Hypothesis (BQH), or weaker interactions and resource specialization. To distinguish between these outcomes, we evolved four species for 44 weeks either alone or together in a toxic pollutant. These species initially facilitated each other, promoting each other’s survival and pollutant degradation. After evolution, two species (*Microbacterium liquefaciens* and *Ochrobactrum anthropi*) that initially relied fully on others to survive continued to do so, with no evidence for increased mutualism. Instead, *Agrobacterium tumefaciens* and *Comamonas testosteroni* (*Ct*) whose ancestors interacted positively, evolved in community to interact more neutrally and grew less well than when they had evolved alone, suggesting that the community limited their adaptation. We detected several gene loss events in *Ct* when evolving with others, but these events did not increase its reliance on other species, contrary to expectations under the BQH. We hypothesize instead that these gene loss events are a consequence of resource specialization. Finally, co-evolved communities degraded the pollutant worse than their ancestors. Together, our results support the evolution of weakened interactions and resource specialization, similar to what has been observed in competitive communities.

## Introduction

How natural and engineered microbial communities function depends on ecological interactions between their member species. As species adapt to one another and to their environment, these interactions may change, and as a consequence, the overall functioning of the community. ^1^ Being able to predict these evolutionary changes may help to intervene and drive a community towards a desirable function. One could imagine, for example, predicting how the gut microbiome would respond to an intervention against inflammatory bowel disease, or how a community in a microbial bioremediation system could be controlled to evolve toward a more stable, efficient state^2–5^.

Evolutionary prediction and control relies on understanding how selection acts on interactions between species. One way to study how these inter-species interactions evolve is to perform experimental evolution by passaging multi-species communities over sequential batch cultures or in chemostats over long time-periods, and following ecological changes in the relative abundances of different species as well as phenotypic and genotypic changes in each community member. Prior studies using this approach have found that microbes can rapidly adapt to both biotic and abiotic factors^6–10^, but being embedded within a community can limit adaption to abiotic factors^8,11–15^. In terms of inter-species interactions, bacterial communities that initially displayed negative interactions evolved towards neutral^9,16^ or positive interactions^8^. This evolutionary response is intuitive, as species can be expected to reduce resource competition and niche overlap^12,17,18^ and may adapt to use resources generated by other species^8,12,16,19^. Accordingly, species evolving in isolation tend to extend their niches in absence of competition and compete when reintroduced into the community context^8,13^.

In contrast, studies that have experimentally evolved communities beginning with positive or facilitative interactions mostly contain only two species or two strains of the same species, often with strong dependencies on one another^10,20–26^. This may be because microbial isolates tend to compete with one another when co-cultured in the lab^27^, meaning that a synthetic community assembled in the lab is unlikely to spontaneously display several positive inter-species interactions. We expect three different outcomes compared to initially competitive communities (Fig. 1): First, if positive interactions are constant and bi-directional over many generations, this might select for each species to increase its positive effect on the other, resulting in mutualism^28,29^. Second, species evolving together might evolve to exploit resources that are provided by others, resulting in stable co-existence because the providing species itself depends on the resource. As proposed by the Black Queen Hypothesis^30,31^, a common consequence of the reliance on public goods produced by others is that the receiving species are selected to lose genes for costly product pathways^21,32^. Third, positive interactions can weaken, particularly if the cooperative traits are costly, resulting in reduced reliance of species on one another^33^. If each species grows independently, one might expect species to evolve to each specialize on a different resource, thereby exploiting available niches more efficiently ^8^.

**Figure 1:**
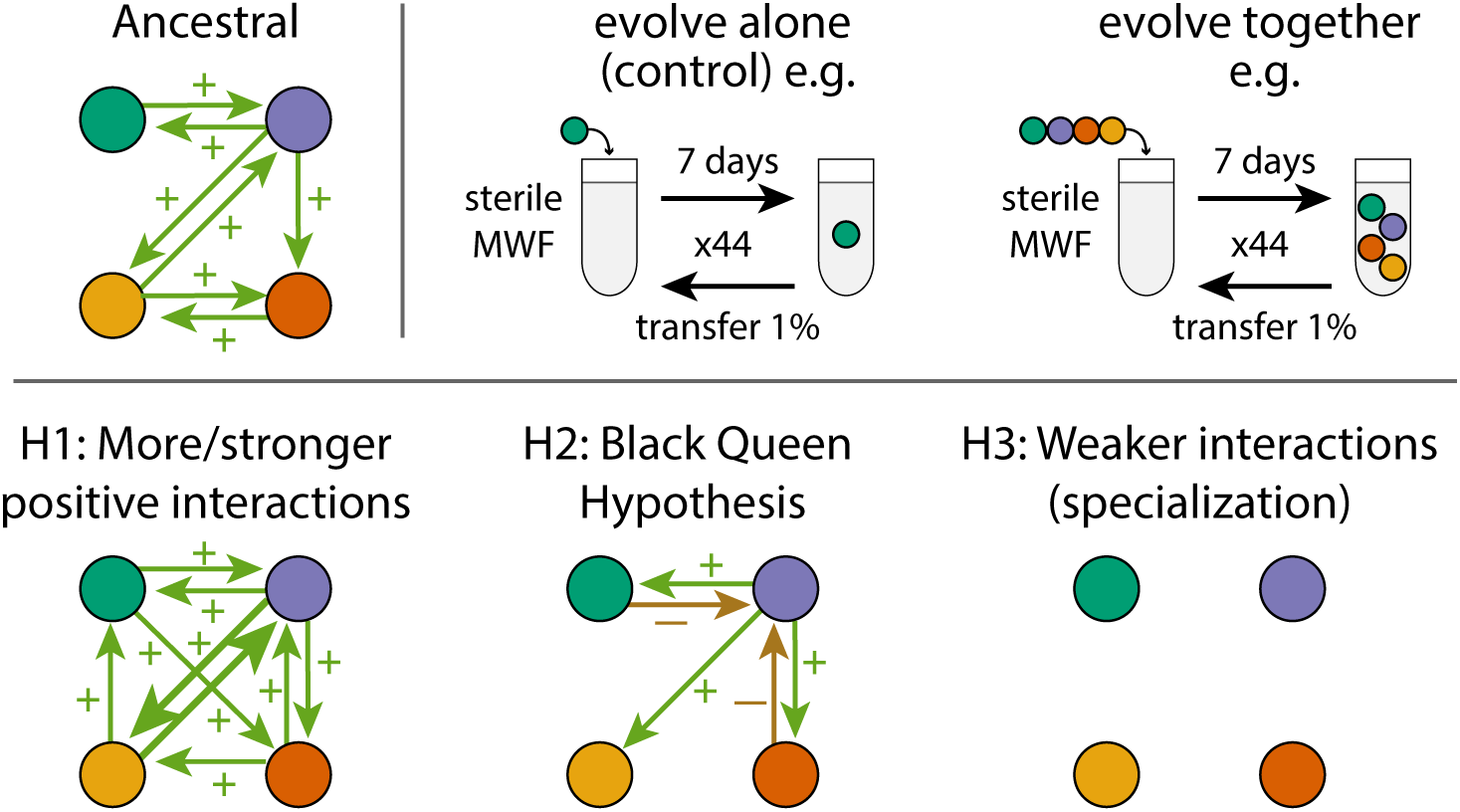
Experiment and hypotheses. Top: An ancestral community with facilitative interactions was evolved in MWF using serial transfers every 7 days for 44 weeks. Different species combinations were explored with species evolving either alone or in a community. Bottom: Hypotheses for how interactions in the community might evolve, assuming that they continue to coexist. H1: More/stronger positive or mutualistic (bi-directional positive) interactions. H2: According to the BQH, few species would provide the public goods for others that would lose the ability to produce them and possibly exploit the producing species. H3: Each species specializes on a different environmental niche, resulting in weaker inter-species interactions.

In our previous work ^34^, we studied a community composed of four bacterial species (*Agrobacterium tumefaciens* (*At*), *Comamonas testosteroni* (*Ct*), *Microbacterium liquefaciens* (*Ml*) and *Ochrobactrum anthropi* (*Oa*)) and showed that facilitation was more prevalent when the community was grown in a toxic environment, in agreement with the Stress Gradient Hypothesis^35^. The toxic environment in question is an emulsion of machine oils used in the manufacturing industry called Metal Working Fluids (MWF), which the four species were capable of degrading when together. They are not known to have a common evolutionary history and were isolated from distinct MWF samples^36,37^. This community represents a tractable model system for exploring how the abiotic and biotic environment shapes the evolution of positive inter-specific interactions and how they relate to community function, in this case, MWF bioremediation.

In this study, the four bacterial species were grown in MWF and left to evolve either in isolation or together in communities (Fig. 1, top) ^8,13^. We quantified bacterial growth and MWF degradation efficiency, and identified genomic changes. By the end of the experiment, positive interactions had declined between the two species evolving together that were able to grow on their own, but not for those that still relied on others to survive and grow. The species evolved in isolation were more productive than those evolved in community and tended to compete with one another when co-cultured. We found little evidence to support the Black Queen Hypothesis, as the species that experienced gene loss events did not increase their reliance on others to grow. Gene loss may instead be a signature of resource specialisation. These results suggest that evolving communities that begin with positive interspecies interactions can evolve similarly to those that begin with negative interactions. In our system, interactions weakened whenever dependencies disappeared, possibly due to niche partitioning, and the evolution of individual species was constrained by coexisting species.

## Results

### Replicate microcosms for each species combination behaved similarly and converged to even communities

Our central question is how facilitative inter-species interactions drive evolution within a microbial community. We addressed this question using experimental evolution of four species either together in groups of 3 or 4 species, or alone as a control. The choice to include this particular 3-species combination was based on preliminary data suggesting that *Oa* may affect community dynamics. While this intuition was confirmed, we do not compare the 3- and 4-species communities explicitly, but nevertheless include all combinations in our data set.

Over the first few weeks, population sizes experienced large fluctuations, which were less pronounced when species were evolving together. When evolving alone, *Ml* and *Oa* went extinct after the first transfer (data not shown), which was unsurprising as they do not grow alone in MWF unless the other species are present ^34^. When evolving alone, *At* only persisted in 2 out of 5 lines (henceforth CAt for “combination” *At*), while *Ct* survived in all 5 microcosms (henceforth CCt). The population sizes of both species dropped initially, but stabilized after about 6 and 11 transfers, respectively (Fig. 2B-C). When species were evolving together, population sizes stabilized after about 4 transfers in the 3-species community (CAtCtMl, Fig. 2D) and 22 transfers in the 4-species community (CAtCtMlOa, Fig. 2E), with the exception of *Ml* that went extinct in 2 out of 5 microcosms in the four-species community.

**Figure 2:**
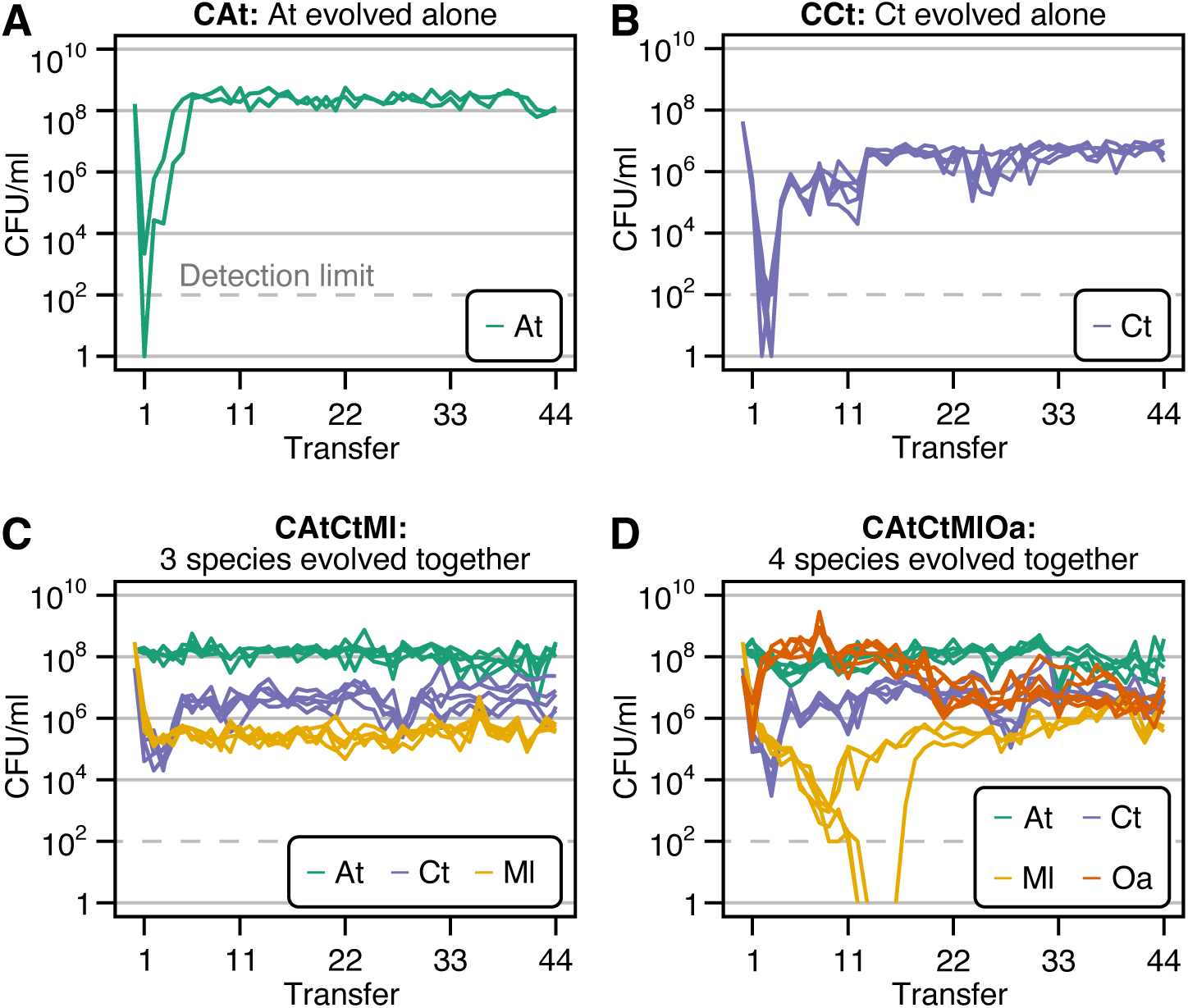
Population sizes over time. Experiments were started with each species in batch mono-culture or co-cultures of three and four species. Every week, we serially transferred cultures by diluting them 100-fold in fresh MWF for 44 weeks. Before each transfer, species abundances were quantified by selective plating. Each species combination (abbreviated as “**C**” followed by the species combination inoculated at the start, e.g. **CAtCtMl**) initially consisted of 5 microcosm replicates (culture tubes). CFU/ml counts from selective plates are shown for all combinations: *At* evolved alone (green), where 3 microcosms dropped below the detection limit (at 10^2^ CFU/ml and is indicated with a grey horizontal line) and were discontinued (A), *Ct* evolved alone (blue) (B), *At*, *Ct* and *Ml* evolved together (C) and *At*, *Ct*, *Ml* and *Oa* evolved together (D). In this species combination, *Ml* dropped below the detection limit in three microcosms, but recovered in one.

In all microcosms where species did not go extinct, the population dynamics in replicate microcosms of the same species combination were similar. By transfer 44, communities were quite even, with relatively small differences in population sizes between species that evolved together (Fig. 2D, E), as expected based on similar studies^16^. The total population sizes in the two co-evolving communities did not increase over time (as observed in^21,38^, e.g.), suggesting that species did not evolve to increase their own or other species’ yield^16^. In fact, fitting a linear model to the total population size in evolving communities (CAtCtMl and CAtCtMlOa) showed a small yet significant decrease over transfers (slope=*−*4.2 *×* 10^6^, P*<* 10*^−^*^9^). Species that evolved alone instead showed no significant change (CAt: P=0.21) or increased over time (CCt: slope=1.4 *×* 10^5^, P*<* 10*^−^*^15^). Species that survived until the end of the experiment went through approximately 300 generations (Table S1).

### Positive species interactions weakened when evolving together

We first explored whether interactions between the evolved species differed from the ancestral ones. We focused on the four species that evolved together (CAtCtMlOa), and to represent the most abundant, genotypically-distinct sub-populations of each evolved bacteria, mixed equal proportions of ten isolates of each species coming from transfer 44 of the same replicate microcosms (Fig. 3A (ii)). We used these mixes, as we detected some within-species phenotypic diversity in growth and degradation (Fig. S1, Fig. S2), but obtained similar results using only one isolate per species, suggesting that growth patterns are consistent across approaches (Fig. S3, Fig. S5). From now on, when referring to *species* in these evolved cultures, we mean these isolate mixes.

**Figure 3:**
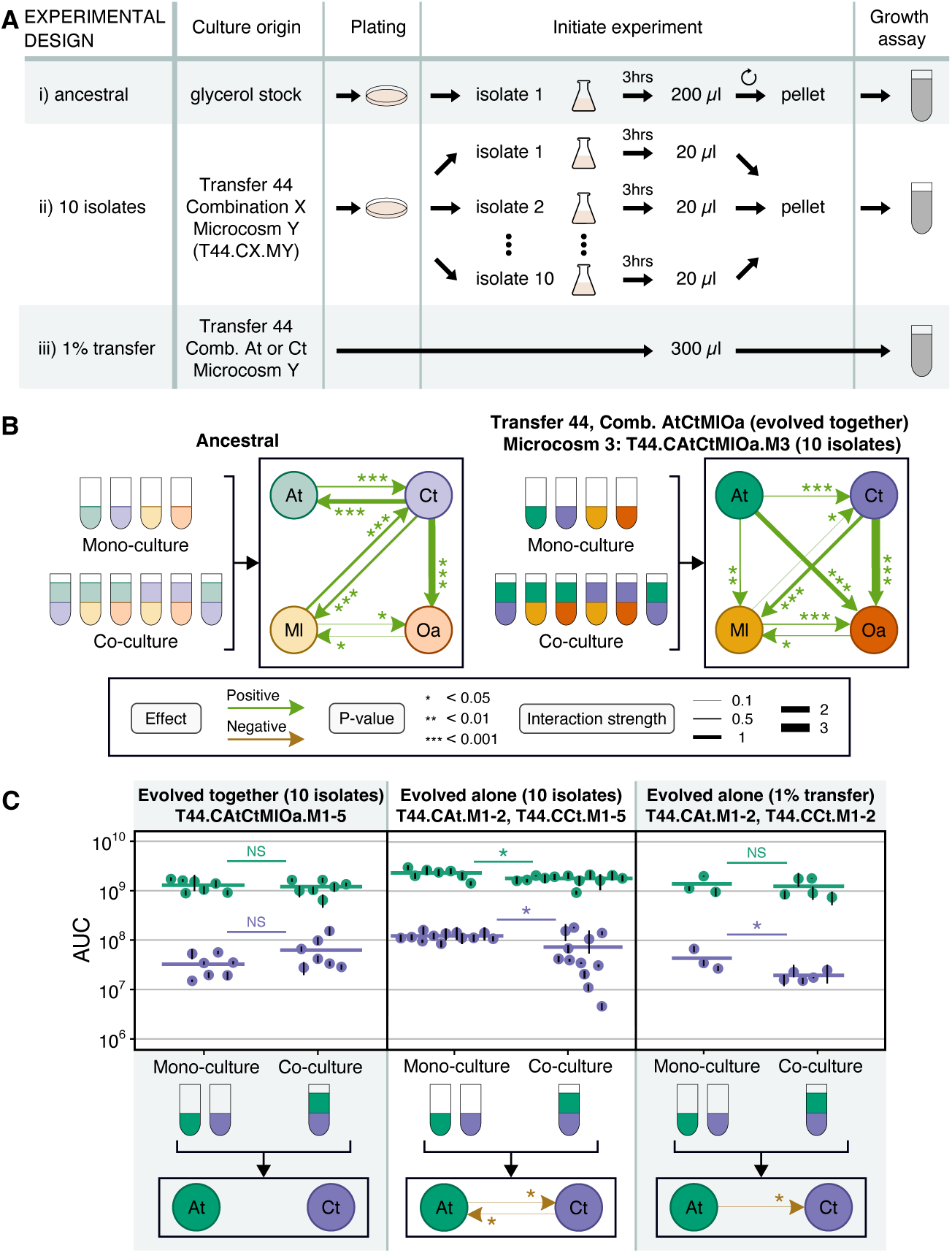
Inter-species interactions. (A) Growth assay experimental design. (i) Glycerol stock of ancestral isolate was grown alone for 3 hours to exponential phase, then washed and resuspended in MWF. (ii) Ten isolates of the same species from transfer 44 of a given species combination and microcosm replicate were randomly picked and grown alone 3 hours to exponential phase, then washed, resuspended and mixed in equal proportions in MWF. (iii) Microcosms from transfer 44 were diluted 100-fold in fresh MWF. We only did this for CAt and CCt (evolved alone), as we couldn’t separate species from one another in CAtCtMl and CAtCtMlOa. (B) Inter-species interaction network in ancestral species (adapted from^34^) versus species evolved together in a community of four for 44 transfers (CAtCtMlOa, microcosm 3) during 12-day growth assays. (C) Interactions based on AUC between *At* and *Ct* evolved together (first column, CAtCtMlOa) or evolved alone (2nd and 3rd column, CAt and CCt, protocols ii and iii from panel A) during 8-day growth assays. For growth assays for CAtCtMlOa (first column) we only co-cultured isolates that had evolved together in the same microcosm, and analyzed all microcosms, with microcosm 3 carried out 3 times (n=7). Matching mono-cultures were done in parallel. For the 2nd column, we mono-cultured CAt.M1 *×*3, CAt.M2 *×*4 (n=7), CCt.M1-2 *×*3 each and CCt.M3-5 *×*2 each (n=12). We co-cultured all possible combinations of microcosms that had evolved alone (CAt.M1 + CCt.M1, CAt.M1 + CCt.M2, etc.) with CAt.M2 + CCt.M1 and CAt.M2 + CCt.M2 carried out twice (n=12). For the 3rd column, we only tested four combinations: CAt.M1 + CCt.M1 (twice), CAt.M1 + CCt.M2, CAt.M2 + CCt.M1 and CAt.M2 + CCt.M2 (n=5). Dots show means and black bars standard deviations of the AUCs, thick horizontal lines show the means of the dots. Significance was calculated using a generalized linear model taking into account microcosm and biological replicate.

Using these isolate mixes, we measured the inter-species interactions in one microcosm where four species evolved together (CAtCtMlOa, replicate microcosm 3, arbitrarily selected among microcosms where all four species were present at transfer 44). We incubated each species in mono-culture or in pairwise co-cultures with each of the other species from the same microcosm over 12 days (Fig. 3B). In mono-culture, contrary to its ancestor (Fig. S4), *At* was able to survive and grow in MWF alone. Both ancestral and evolved *Ct* were able to survive and grow in MWF (Fig. S3C), but the area under the growth curve (AUC) of evolved *Ct* was significantly lower across different assays (Fig. S5, Fig. S6B). Finally, *Ml* and *Oa* from all microcosms were still unable to grow alone (Fig. S8).

By comparing the AUCs of mono-cultures with pair-wise co-cultures, we were able to reconstruct an interaction network (Fig. 3B), as previously done for the ancestral network in Piccardi et al. ^34^ . *Ml* and *Oa* continued to rely on *Ct* for survival, but we found no evidence for increased mutualism between *Ml* and *Ct* (Fig. S9). Unlike the ancestral community, evolved *At* promoted the survival and growth of *Ml* and *Oa*, while it no longer benefited from evolved *Ct*. The appearance of positive interactions towards the two species that could not grow alone was expected because *At* could now grow independently ^34^. The lack of competition between the two independent species (*At* and *Ct*) was however unexpected, as our intuition from previous work was that autonomous species should compete^34^. This motivated us to explore this relationship further.

To understand whether the weakened interaction between *At* and *Ct* was consistent across all five microcosms where the species had evolved together (CAtCtMlOa), we compared the growth of evolved *At* and *Ct* isolates from the same microcosms in mono- and pair-wise co-cultures. We found that the AUC, the maximal CFU/mL difference between two consecutive days of each species (a proxy for growth rate), and the maximum population size of *At*, did not differ significantly when co-cultured with *Ct* from the same evolved microcosm (linear model with biological replicate as a random effect; AUC: *P* = 0.65, Fig. 3C, left column; maximal CFU/mL difference between two consecutive days: *P* = 0.37, Fig. S7B, left column; maximum population size: *P* = 0.56, Fig. S7C, left column). Instead, *Ct* had a significantly greater maximal growth rate when co-cultured with *At* from the same evolved microcosm, but its AUC and its maximum population size did not differ significantly (linear model with biological replicate as a random effect; AUC: *P* = 0.1275, Fig. 3C, left column; maximal CFU/mL difference between two consecutive days: *P* = 0.0265, Fig. S7B, left column; maximum population size: *P* = 0.123, Fig. S7C, left column). In other words, taking into account all microcosms and several ways to measure interactions, *At* and *Ct* no longer interacted significantly.

### Species that evolved alone tended to interact negatively

We wondered whether the reduction in positive interactions between *At* and *Ct* when evolved together was simply the result of adaptation to the harsh MWF conditions. We compared the growth of *At* and *Ct* that had evolved alone when grown in mono- and pair-wise co-cultures (Fig. 3C, middle column). Both species inhibited each other’s growth, where the AUC (linear model with biological replicate as a random effect, *Ct → At P* = 0.015, *At → Ct P* = 0.01) and maximal population size (*Ct → At P* = 0.004, *At → Ct P* = 0.049) (Fig. S7A, C) of the co-cultures were lower than the mono-cultures. Although the effect sizes do not appear large on the plot, they are non-negligible (e.g. AUCs of *At* and *Ct* were reduced by 22.8% and 40.5% on average, respectively). Overall, this suggests that the evolutionary response of *At* and *Ct* is different whether they evolve alone or in the community context.

One explanation for the competitive interactions may be that the isolates we used for these assays had a particularly high fitness within their populations. To test whether our results were biased in this way, we transferred the entire populations of *At* and *Ct* from two microcosms each where they had evolved alone directly into mono- or co-culture assays (Fig. 3A (iii)). *At* still inhibited the growth of *Ct* (linear model with biological replicate as a random effect, At *→* Ct *P* = 0.045) (Fig. 3C, right column), suggesting that there was likely nothing particular about the 10 isolates. In sum, the positive interactions between *At* and *Ct* in the ancestral strains switched toward more neutral interaction when evolving together, and competition when evolving alone.

### Species evolved alone were more productive than those evolved together

A possible explanation for why species that evolved alone compete with one another in coculture, is that evolving alone allows them to increase their niche coverage, resulting in competition with future invaders into its environment. If instead, a focal species is already sharing the environment with other species with which it does not compete, their presence may prevent the focal species from expanding its niche thereby limiting competitive interactions from arising over evolutionary time-scales. While niche partitioning is difficult to quantify in a complex chemical environment like MWF, we predicted that if species that evolved alone cover more niche space, they should grow faster or to a larger population size compared to their counterparts that evolved with others. Consistent with this prediction, the AUC of *At* and *Ct* that had evolved alone was significantly higher than their counterparts that had evolved in community (linear model with biological replicate as a random effect, *At P* = 0.001, *Ct P <* 0.001), even when they were grown in co-culture (linear model with biological replicate as a random effect, *At P <* 0.001, *Ct P <* 0.0036, Fig. 3C, S1, S10). While these results do not prove that evolving alone led to greater niche expansion (they may simply have evolved higher yield), they match observations from previous studies^8,9,12^ showing that adaptations to increase productivity are limited when species are evolving with others.

### Ecological context influences genomic changes

Given the differences between *Ct* and *At* that had evolved alone or together, we next wondered whether we could find corresponding genomic variations and determine when they emerged. To this end, we extracted and sequenced the DNA of all microcosm populations every 11 transfers and reconstructed their evolutionary trajectories (see Methods). Because we lack statistical power for *At* (it only survived in 2 microcosms when evolving alone), we focused on *Ct*.

We observed distinct patterns for *Ct* evolved alone or together with other species (Fig. 4). When evolved with other species (CAtCtMl and CAtCtMlOa), *Ct* accumulated a higher number of variants compared to when it was evolving alone (CCt, Kruskal-Wallis chi-squared = 6.818, P = 0.009, Fig. 4B left), resulting in a higher total allele frequency (Fig. S13). But many variants did not fix and remained at intermediate frequencies (Fig. 4A center and right). Instead, when evolved alone, a significantly higher number and proportion of variants fixed (number: Kruskal-Wallis chi-squared = 4.165, P = 0.041; proportion: Kruskal-Wallis chi-squared = 4.810, P = 0.028, Fig. 4B center). This suggests suggests hard sweeps when evolving alone and soft sweeps when evolving in community, which can be explained by the strong drop in population size early on in the experiment when alone compared to in community (Fig. 2B versus C and D).

**Figure 4:**
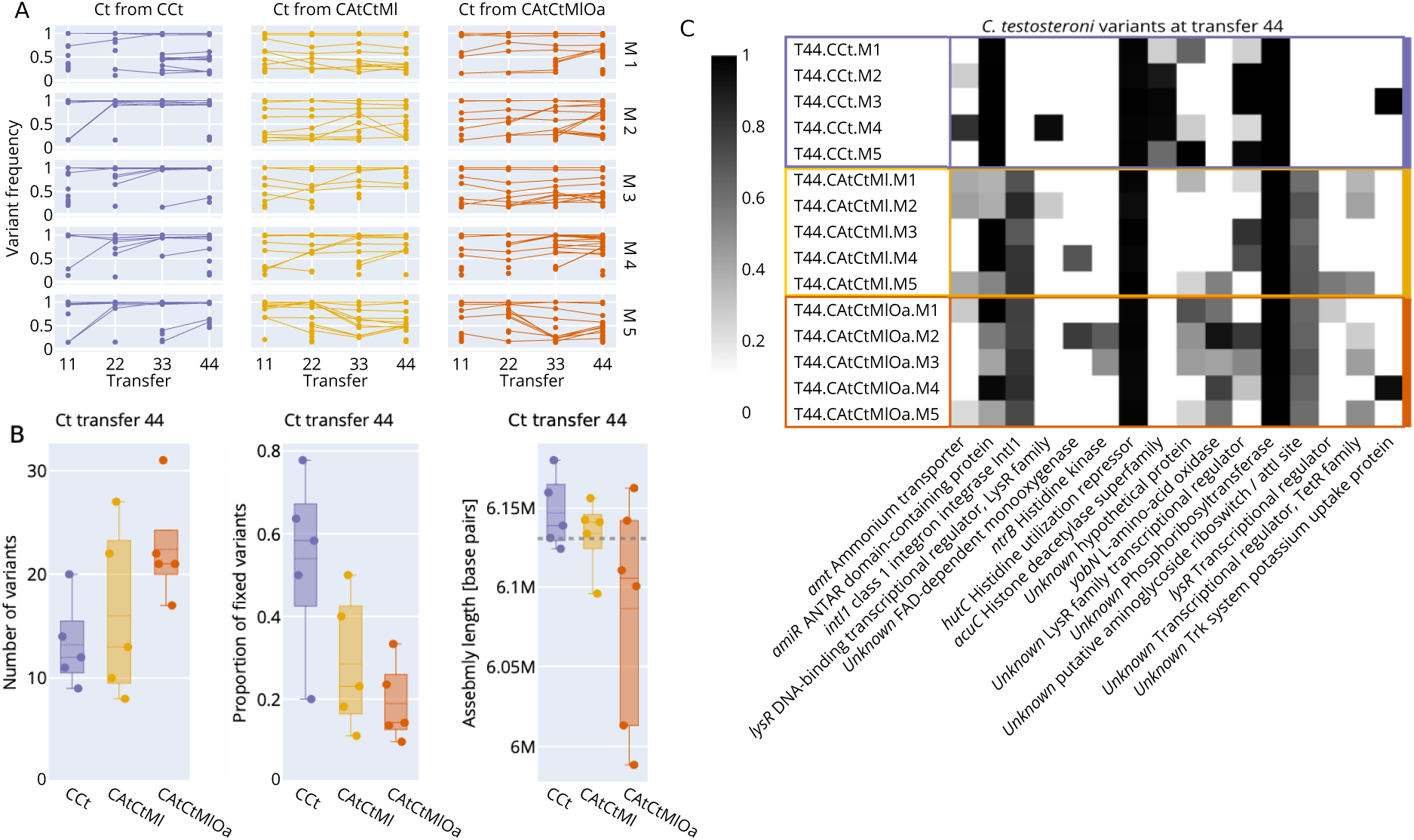
Genomic changes. (A) Variant frequency trajectories in all *Ct* populations. Each dot/line represents a different variant at a different location. (B) Number of variants found in each *Ct* population at transfer 44 (left, matches data in panel A), proportion of variants that reached fixation (center, matches data in panel A), and de-novo long-read assembly lengths based on PacBio sequencing of selected isolates from transfer 44 (right). The dashed line represents the assembly length of the ancestor. (C) Mutated genes with protein annotation that were found in at least two *Ct* populations. The grey shade indicates the frequency of the mutated allele.

Given that ecological context affected allele frequencies and fixation rates, we expected variant targets to also depend on the presence or absence of other community members. We annotated the variants and filtered for genes that were mutated in at least 2 microcosms (Fig. 4C). One gene (*acuC*), which codes for a histone deacetylase, was mutated exclusively in CCt (in all 5 microcosms). Mutations to seven genes were exclusive to combinations CAtCtMl and CAtCtMlOa (2 genes were affected in all 10 microcosms), and two genes were mutated and almost completely fixed in all microcosms across all species combinations, likely related to adaptation to MWF.

In *Ct* coming from one particular microcosm (T44.CAtCtMlOa,M2), we observed that 3 out of 10 isolates were able to grow alone at two-fold higher MWF concentrations and had no measurable lag time, while the remaining 7 grew more characteristically for this species (Fig. S1, S2). We whole-genome sequenced one isolate from each subpopulation and identified a mutation in *ntrB* (coding for a Histidine kinase) in the more resistant strain. We confirmed that this variant was present but not fixed in the metagenomic sequencing data of that population. The resistant isolate also had a large deletion (Fig. S15C), which we discuss below. Why this more resistant variant did not fix, and whether the wildtype-like variant is acting as a cheater is unclear.

### No evidence for Black Queen dynamics in our system

The Black Queen Hypothesis^30,31^ (BQH) predicts that if several species in a community are contributing to a public good, all but one species should lose this trait, leading to gene loss in evolving communities. In our system, environmental detoxification can act as a public good. Although we do not know which genes are involved, we explored whether gene loss occurred preferentially for species evolved together compared to alone by long-read sequencing whole genomes of isolates from all microcosms at transfer 44 (see Methods). After assembling full *Ct* and *At* genomes, we found that two *Ct* isolates from CAtCtMlOa were over a 100k base pairs shorter than the reference genome. We mapped these to the reference strain and found an identical deletion of 145k base pairs including 31 genes (see Fig. S15). We doubt that these deletions are due to increased dependence on other species in the community, as the BQH would predict, as these two isolates grew similarly in isolation to the ones without the deletion (Fig. S1). Indeed, one of these isolates was the strain that was resistant to higher MWF concentrations described above and grew better than the other isolates (Fig. S1, S2). For *At* we observed a large deletion in one isolate from CAtCtMl, but nothing striking for *Ml* and *Oa* (Fig. S13B). Despite these observations, we lack statistical power to conclude anything general. We also used the assemblies to check if any sequences from other species were integrated in the genomes, however, no transfer events were detected.

As it seemed plausible that 44 weeks were too short for structural changes to occur systematically, we next explored whether point mutations in regulatory regions might have instead led to down-regulation in gene expression in evolved communities. We extracted and sequenced RNA from isolates of all microcosms of *Ct* and *At* at transfer 44 as well as their ancestors. As the quality of RNA from *At* samples was low, we focused on *Ct*. Contrary to the prediction of the BQH, we found no significant difference in the normalized expression levels from the isolates of CAtCtMl and CAtCtMlOa that had evolved in community, while several genes in isolates that had evolved alone (CCt) were significantly down-regulated when compared to the ancestor (Fig. S13C-E, Table S2). The only mutation present uniquely and repeatedly in CCt was in the *acuC* gene, which is expected to affect gene expression.

### *At* degrades MWF better after evolving alone but not in community

Next, we investigated whether the decline in positive inter-specific interactions over the 44 transfers was associated with a shift in MWF degradation efficiency (as in Rivett et al. ^16^). If, for example, co-evolved species have indeed reduced their niche overlap and diverged in their resource use, we might expect greater overall MWF degradation. On the other hand, *Ct* that evolved in the community grew slower than its ancestor, which may lead to worse degradation, as it is one of the main degraders in the community^34^.

Over the 44 transfers (Fig. 5A), *Ct* and the two evolved communities reduced their degradation efficiency, such that at the end, they degraded less than their ancestral counterparts (%COD on day 7, isolates from transfer 1 vs. 44 in *Ct* evolved alone, CAtCtMl and CAtCtMlOa, respectively: paired t-tests, *t* = *−*5.7165, *P <* 0.01; *t* = *−*14.641, *P <* 0.001; *t* = *−*18.131, *P <* 10*^−^*^4^). In contrast, the two microcosms in which *At* evolved alone degraded significantly better than their ancestral counterpart (%COD on day 7, isolates from transfer 1 vs. 44 in mono-evolved *At*: linear model, *t* = *−*20.91, *P <* 10*^−^*^5^, Fig. 5A) and even compared to all other microcosms (%COD on transfer 44, day 7, comparing *At* with *Ct* and the 3- and 4-species communities, respectively: linear model, *t* = 10.85, *P <* 0.001; linear model, *t* = 10.35, *P <* 0.001; linear model, *t* = 8.274, *P <* 0.001, Fig. 5A).

**Figure 5:**
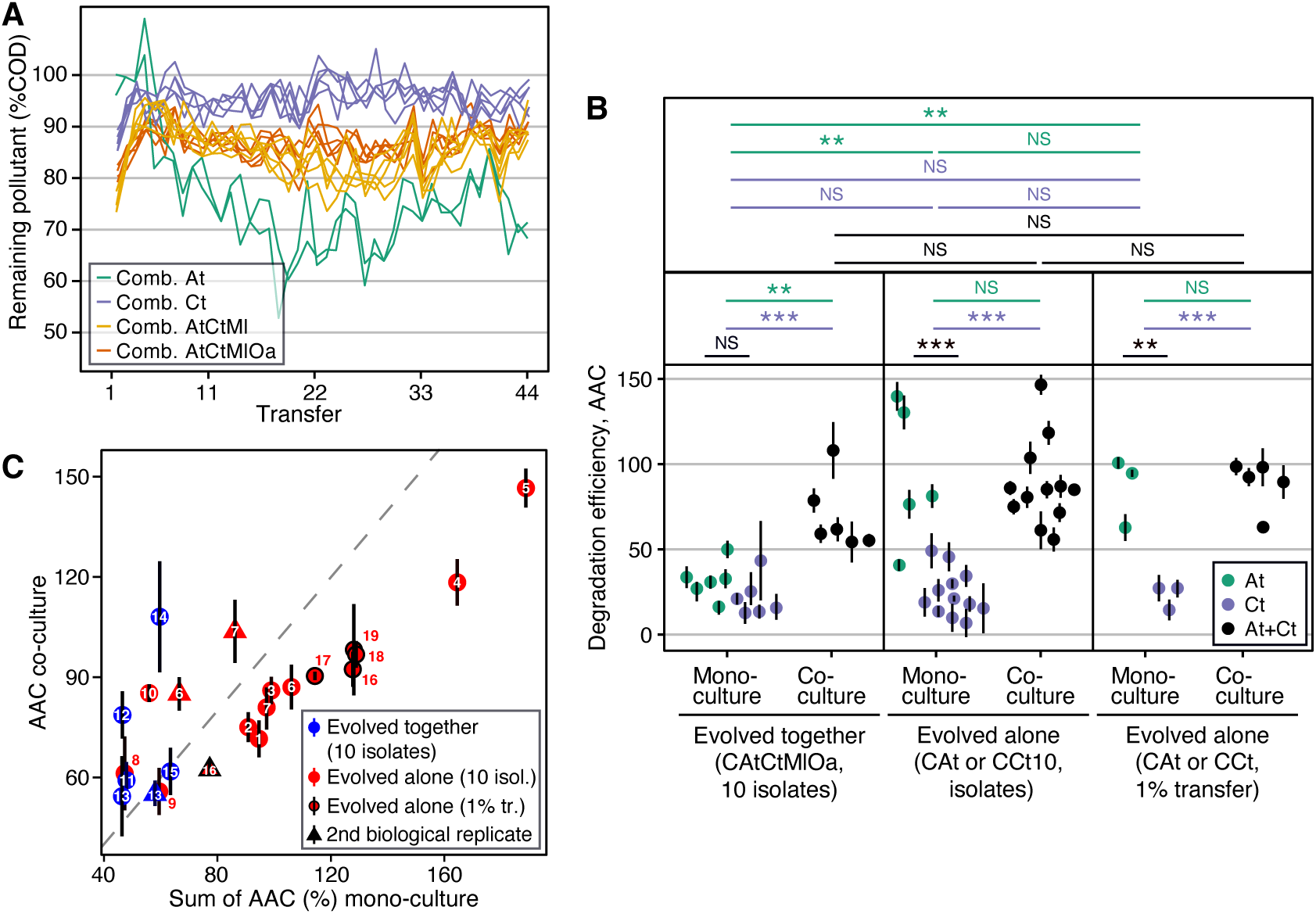
Degradation efficiency. (A) Remaining pollutant, measured as chemical oxygen demand (COD, g/L) as a percentage of the COD of an abiotic control for each microcosm before each transfer in all four species combinations (lower implies greater degradation). (B) Comparison of the degradation efficiency of *At* and *Ct* ancestral, evolved alone or together with others, where the Area Above the degradation Curve over 8 days is shown (higher AAC implies greater degradation, unit is sum of percentages). (C) Prediction of an additive model of the sum of degradation efficiencies of individual species is plotted against degradation efficiency of the co-cultures of mono-versus co-evolved species. Points lying above (or below) the dashed line degrade more (or less) efficiently than predicted by the additive model. Each small number associated with a datapoint indicates a given species combination (e.g. CAt.M1 + CCt.M1). Certain numbers have both a circle and a triangle, which are biological replicates of the same species combination. A list of which number corresponds to which combination can be found in Dataset 1.

Knowing that *At* was a member of the two evolved communities, we wondered why the degradation efficiency of the communities was worse than *At* evolved alone. Did the community members inhibit the degradation efficiency of *At* or did it not evolve improved degradation? We find evidence to support the latter: when grown alone, *At* from the evolved community degraded less efficiently than when it had evolved alone (%AAC, assays with 10 isolates and 1% transfer, respectively: linear model with biological replicate as random factor, *t* = 3.590, *P <* 0.01; *t* = 5.373, *P <* 0.01, Fig. 5B). This mirrors our earlier observation that *At* that had evolved alone grew to greater population sizes than when evolved in community (Fig. 3C, S1, S10, compare *At* mono-cultures). These data suggest that other species may have constrained the evolution of *At*, preventing it from evolving greater degradation efficiency by occupying some niches that it could instead fill when evolving alone. If the species that evolved together with *At* are filling the available niches, might they complement *At*’s ability to degrade MWF?

### Species evolved together degrade MWF synergistically

Following our observation that *At* evolved in community does not degrade as much as when it evolved alone, we wondered whether the species evolving together with *At* – notably *Ct* – could improve its degradation efficiency. By applying an additive null model to degradation efficiency, we compared the combined degradation of the two mono-cultures of these two species to degradation in their corresponding co-cultures^9,16,34,39^ (Fig. 5C). Although there were some differences between experimental repeats with different sub-samples of the evolved populations, overall, we found that *At* and *Ct* that had evolved in the same microcosms had small positive effects on each other’s degradation efficiency (statistical analysis in Fig. S11). For the species that had evolved alone, depending on which isolates we used for the assays, we found that in some cases the two species significantly reduced each other’s degradation efficiency (Fig. S11). Together, this supports the hypothesis that co-evolving species do not overlap much in their niches and can therefore synergistically degrade MWF. Instead, *Ct* that had evolved alone seems to interfere with the degradation ability of *At* that had evolved alone, which suggests that the potential of *At* to expand its niches and increase its own degradation efficiency may have been limited when it evolved in the community context.

## Discussion

Our main goal was to establish how interactions within a facilitative community might change as species evolved together. Would interactions become more mutualistic, would one species becomes parasitized by the others to produce all the public goods (as per the BQH) or would they evolve to specialize or even compete (Fig. 1)? Similar experiments with initially competitive communities found that interactions weaken as their members co-evolve to specialize on different resources^6,8,9,13,16,40^. In our 44-week long evolution experiment, species that relied heavily on others to survive in MWF continued to do so, but with no evidence of strengthened mutualism. Instead, one species *At*, that evolved to grow independently in MWF weakened its positive interaction with the other independent species, *Ct*. When we evolved each of those two species alone, they competed when put back together.

Our interpretation of this outcome – while well aware that alternative explanations exist – is that in the community, *At* and *Ct* experienced weak selection to expand into occupied niches and compete with other residents, driving them to specialize on more available resources (H3 in Fig. 1), analogous to character displacement in Darwin’s finches^41–43^. Instead, when evolving alone, they may have become generalists by expanding into available niches because no other species were occupying them^12^. The presence of other species may then have constrained the evolutionary potential of *At* and *Ct* (similar to results reported by Hall et al. ^12^). Evidence for this is that after evolving alone, isolates of these two species grew significantly better than those that had evolved in community. Indeed, *At* evolving alone was the only condition where degradation improved over the course of the experiment and largely surpassed the degradation ability of the community, even though the community includes *At*. In follow-up experiments reported elsewhere we also found that new, non-resident species were more likely to invade the ancestral compared to the evolved community^44^, suggesting that the community members evolved to cover the available niche space. An alternative initial hypothesis was that positive interactions might increase, leading to the evolution of mutualism (H1 in Fig. 1) because mutants that overproduce public goods should be favored as they promote the growth of species that “help” them^28,29^. This outcome can result in increased community productivity^22,45^, increased aggregation between cooperating strains^20,23,25^ and/or loss of independent growth ^21^. By comparing the ancestral and evolved interaction networks (Fig. 3), the bi-directional interactions between *Ct* and *Ml* were a candidate for this. However, we found no significant increase in the strength of their positive interactions, at least in this one microcosm (Fig. S9). Second, the number of positive interactions may increase if species generate new niches, which others can evolve to occupy^8,24,46^. While it may appear that there are more positive interactions in the evolved community (Fig. 3), the positive effects of *At* on *Ml* and *Oa* were already observed in the ancestors growing under conditions where *At* survives^34^, and are not newly evolved traits. In addition, if stronger positive interactions had evolved, we would expect overall community productivity to go up because resource use becomes more efficient ^8,38^. While we do find some synergy in MWF degradation, the co-evolved co-cultures still degrade less than *At* evolved alone, and total population sizes even decreased over the evolutionary experiment.

The other question was whether we would find support for the Black Queen Hypothesis^30^ (H2 in Fig. 1): if several species in the ancestral community provide a “service”, others should evolve to lose it, manifesting itself in gene loss for species evolving together^30,47^. What constitutes a “service” in our context is not clear mechanistically, but *At* and *Ct* do facilitate the other two species by detoxifying the environment ^34^. If they were initially achieving this in overlapping ways, the two species might evolve to specialize on degrading different toxins. This would predict greater gene loss or reduced gene expression in the species evolved together compared to those evolved alone, and greater reliance on one another for survival. We found little evidence in support of this prediction: two *Ct* isolates that had evolved in community experienced large deletions, but these isolates grew similarly alone to others without the deletion, and at least one of them was even more resistant to MWF compared to a strain that had evolved alone.

These findings made us realize that the evolution of resource specialization within a community predicts similar patterns of gene loss to the BQH, as the ability to use certain resources that are already taken up by others becomes superfluous (Fig. S16). Given that our data generally support niche specialization rather than increased reliance on other species (at least for the two species we focused on), the deletions we see may be more in line with specialization rather than the BQH, but additional work would be needed to test this idea. In other words, we suggest that the BQH and specialization are two similar processes that we expect to drive genomic changes when species evolve in community. By itself then, gene loss alone should not be taken as evidence supporting the BQH.

Why the bacteria evolved to degrade less in all conditions except for *At* evolving alone, is an important open question when optimizing microbial community function. One possibility is that in the communities and when *Ct* was alone, selection favored the emergence of cheaters that grew faster but contributed less to MWF degradation, which may have increased the death rate, explaining the lack of increase in total population size^48,49^. Alternatively, cells might have evolved to resist the toxins without secreting toxin-degrading enzymes, for example by thickening the cell wall or using efflux pumps^50–52^. This would make resistance into a “private good” and reduce MWF degradation. Third, the community constrained the evolution of *At*, explaining why its degradation did not improve when evolving in the community. Regardless of the mechanism, our results suggest that the problem of loss of community function needs to be addressed in future studies. Otherwise, single species like *At* might be better suited compared to communities, at least for this particular function of MWF degradation.

A final interesting question in community evolution concerns predictability: Do parallel microcosms evolving under the same condition resemble one another? Previous evolutionary experiments using communities found bimodal or trimodal outcomes in final relative abundances^53,54^. We observed striking parallel ecological dynamics between microcosms, whereby relative abundances converged by week 44, despite the occasional extinction of *Ml*. *Oa* appeared to play a destabilizing role, as population sizes of all species fluctuated more strongly in CAtCtMlOa compared to CAtCtMl before converging (Fig. 2C, D). As in other such experiments^10,20,26^, we also observed some parallelism in genomic evolution, where several mutations and deletions occurred in parallel lines of the same experimental condition, at least in *Ct*. While it is tempting to speculate on the effects these mutations might have, we prefer to leave mechanistic analyses to future work where we would build the appropriate mutants.

One of the weaknesses of our system is that chemical analysis is challenging, meaning that we lack a mechanistic understanding of pollutant degradation or the interactions between species. We are therefore blind to how resources are being partitioned, what lies behind the positive interactions, the consequences of genomic changes, or why degradation efficiency dropped over time in evolving communities. Another difficulty was our inability to generalize, as the community only includes four species, and each followed a different evolutionary trajectory. Running similar experiments using communities with more species in a simpler chemical environment could help to test our hypotheses further.

Our experiments present a case study of how four species can evolve in a toxic environment, showing that for species pairs whose dependencies were facultative, interactions weakened over time. Positively interacting species are therefore not necessarily expected to evolve towards mutualism^29^, and can instead evolve similarly to competitive communities. From an applied perspective, community function dropped over time as the species evolved, suggesting that to maintain function, new strategies are needed. Finally, parallels can be drawn to evolution in other toxic environments, such as those containing antibiotics, a phenomenon that has classically been studied in single species in isolation^55^. Being able to predict and control the evolution of microbial communities would be impactful in many such contexts.

## Methods and Materials

### Bacterial species and culture conditions

The ancestral species used in this study were *Agrobacterium tumefaciens* str. MWF001, *Comamonas testosteroni* str. MWF001, *Microbacterium liquefaciens* str. MWF001, and *Ochrobactrum anthropi* str. MWF001. More details on these strains can be found in Piccardi et al. ^34^ and their genome sequences on NCBI (Accession: PRJNA991498). Note that *Microbacterium liquefaciens* was previously referred to as *Microbacterium saperdae* but a more recent classification has led us to refer to it differently.

All experiments were performed in 30ml batch cultures in glass tubes containing 0.5% (v/v) Castrol Hysol^TM^ XF MWF (acquired in 2016) diluted in water with added salts and metal traces (see Piccardi et al. ^34^ for detailed recipe). Cultures were incubated at 28*^◦^*C, shaken at 200 rpm.

### Evolution experiment

All the experiments (initially 6 treatments: 4 mono-cultures, 1 3-species co-culture, 1 4-species co-culture) were conducted simultaneously in 5 microcosm replicates to give 30 experimental cultures in addition to 3 sterile controls (Fig. 6).

**Figure 6:**
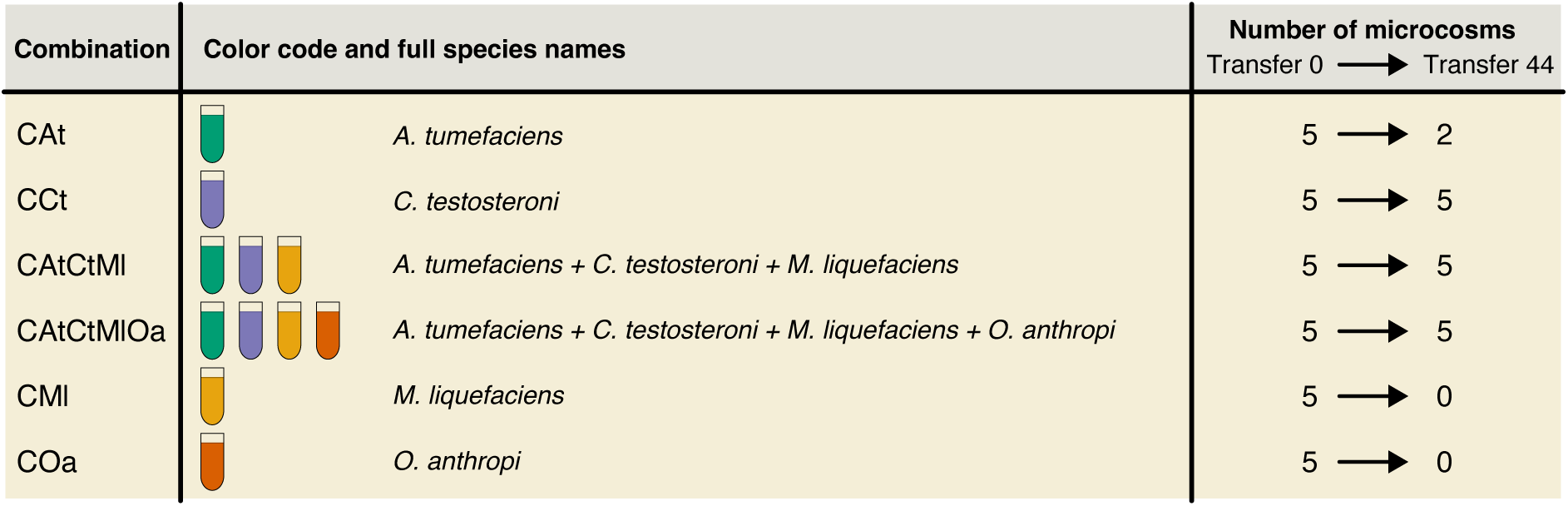
Evolved species combinations.

All tubes were incubated at 28*^◦^*C, shaken at 200 rpm for a total of 7 days. Each week for a total of 44 weeks, 29.7 mL of fresh MWF medium was prepared and 300 *µ*L of the week-old culture transferred into it. Before each transfer, population sizes (CFU/mL) were quantified using serial dilution and selective plating and CODs (pollution load) were quantified using Macherey Nagel 15 g/L COD tube tests (see Piccardi et al. ^34^ for detailed recipe). A sterile tube containing MWF but no bacteria was always used as a control for the COD measurement. Every week, 1mL of the bacterial cultures was harvested for each treatment, spun down at 10,000 rcf for 5 minutes, resuspended in glycerol 25% (diluted in PBS) and stocked at -80*^◦^*C for future analyses (e.g. DNA extraction). All 5 replicate populations of *M. liquifaciens*, *O. anthropi* and 3 replicate populations of *A. tumefaciens* in mono-culture went extinct, and these microcosms were discarded after 10 weeks.

At the end of the experiments (after transfer 44), we collected 10 individual isolates of each species from each population for further analysis by plating populations on selective media and randomly picking 10 colonies. These colonies were then grown overnight in TSB at 28*^◦^*C, shaken at 200 rpm, spun down at 10,000 rcf for 5 minutes, resuspended in glycerol 25% (diluted in PBS) and stocked at -80*^◦^*C.

### Bioinformatic analysis

#### Ancestral lineage sequencing and annotation

DNA coming from each ancestral species was sequenced using a combination of Illumina (MiSeq) and PacBio (RSII). PacBio raw data for each genome sequencing was assembled using canu v. 2.2^56^ and polished with racon v.1.5.0. ^57^ The assembly was further corrected using the Illumina data with polypolish v. 0.5.0.^58^ The assemblies were then annotated using bakta v. 1.2.4. ^59^

#### DNA extraction and sequencing

To extract DNA from the frozen populations for Illumina sequencing, we defrosted the populations from the T-1 transfer (e.g. to sequence transfer 22, we defrosted transfer 21), washed and resuspended the cells in 1ml of PBS and inoculated 300 *µ*L into 29.7 mL of fresh MWF medium. After 1 week, we collected 15mL of each sample, split into 1.5mL Eppendorf tubes and spun down at 10’000rpm for 10 minutes. The bi-phasic supernatant was carefully discarded. Pellets coming from the same sample were resuspended in PBS and pooled together into one single 1.5mL Eppendorf tube. Cells were precipitated and resuspended in PBS twice, to remove any remaining MWF. A negative control was included in the process and followed the same procedure as the samples. To extract DNA from isolates for PacBio sequencing, we grew the previously frozen isolates overnight in TSB at 28*^◦^*C, shaken at 200 rpm, and spun them down at 10,000 rcf for 5 minutes.

The resulting pelleted cells were incubated in 150 *µ*l of lysozyme solution for 30 minutes at 37*^◦^*C. After this incubation period, 5 *µ*l of RNAse solution (5mg/ml) was added. The RNAse treatment was performed for 30 more minutes at the same temperature. The lysozyme action creates pores in the cell wall of the cells, allowing the RNAse to degrade any possible remaining RNA in the sample. After this second incubation period, 600*µ*l of lysis buffer was added to the sample. The lysis buffer solution contains 9.34mL of TE buffer (PH 8), 600*µ*l of SDS 10%, 60*µ*l of Proteinase K and 2*µ*l of ß-mercaptoethanol. Cell lysis was performed for 1 hour at 56*^◦^*C. Once the cell suspension became transparent, 700*µ*l (1v/v) of Phenol-Chlorophorm-Isoamylalcohol (PCI, 25:24:1) was added to the tube. Samples were mixed by inversion for 1 minute and left to rest on ice to allow phase separation. After the phases were clearly visible, the sample was centrifuged at 13’000 rpm for 15 minutes at 4*^◦^*C. The resulting clear supernatant was transferred to a new tube (600*µ*l of volume). PCI cleaning was performed one more time to purify the DNA, resulting in around 500*µ*l of clear liquid containing the suspended DNA. After the DNA cleaning, 50*µ*l of sodium acetate (5M) and 500*µ*l of Isopropnol were added to the sample, allowing the DNA to precipitate. Insoluble DNA was incubated at -80*^◦^*C for two hours and centrifuged down at 13’000 rpm for 15 minutes. The alcoholic supernatant was discarded. The precipitated DNA was washed with 1ml of ethanol 70% (v/v), re-centrifuged at 13’000 rpm for 15 more minutes, and the supernatant removed. The air dried pellet was then redissolved in 50*µ*l nuclease-free water, and the concentration and purity were analyzed using Qubit and Nanodrop.

The obtained DNA was sequenced with using the Illumina platform with two different platforms at the Oxford genomics facilities: Samples from transfer 22 were sequenced using HiSeq4000. While transfers 11,33,44 were sequenced using NovaSeq. The reason behind the different platform usage was the discontinuation of the former at the selected facility. PacBio sequencing was performed on individual isolates of each species from transfer 44 (Table x) at the Lausanne Genomic Technologies Facility using a Sequel II system (SMRT cell 8M).

#### RNA extraction and sequencing

We grew the previously frozen isolates from transfer 44 (see above) overnight in TSB, washed them in PBS and then inoculated 300 *µ*L into 29.7 mL of fresh MWF medium. After 7 days of growth, the cells were pelleted and the RNA extracted using the RNeasy PowerSoil Total RNA Kit. The extraction yielded a minimum of 30 ng/*µ*l in 10 *µ*l. The sequencing library was prepared including ribosomal RNA depletion using the Illumina ZeroPlus library perparation kit and sequenced on a NovaSeq 600 sequencer.

#### Sequence data processing and analysis

For each Illumina sequencing data-set, an initial quality control was performed using FastQC, to evaluate the overall per-position quality, the k-mer enrichment (which could indicate adapter contamination), and the GC-content (which could indicate origin admixture). ^60^ Adapters and low quality sequences were removed using trimmomatic v. 0.36, using the parameters PE, leading=3, trailing=3, slidingwindow=4:15, minlen=60.^61^ The resulting cleaned reads were mapped against the ancestral genome references using minimap2 v. 2.22. ^62^ For sequencing data derived from microcosms with multiple species, the reads were aligned against all merged ancestral reference genomes with no secondary mapping in order to avoid cross-mapping. The mapping was filtered to remove distant alignments and low quality alignments using samtools view with the parameters -f 3 and -q 60.^63^ Based on the filtered alignment files, we identified variants with freebayes version 1.3.6 with the parameters–min-alternate-count 3 –min-alternate-fraction 0.05 –pooled-continuous –haplotype-length 0–standard-filters.^64^ Variants outputted by freebayes were then filtered by a minimum population frequency of 10% and a minimum Phred quality of 20. A variant was considered fixed if it exceeded a frequency of 95%.

PacBio whole-genome sequencing data were assembled using canu version 2.2. ^56^ The resulting assemblies were polished with racon 1.5.0, ^57^ and annotated with bakta v. 1.2.4. ^59^ To investigate potential intra-species gene transfers, we split the assemblies into 150-mers and taxanomically classified the 150-mers using using krakken 2.1.2. ^57,65^ RNA sequencing data was analyzed using the RASflow workflow with default parameters wrapping hisat2 2.1.0 as an aligner, htseq-count 0.11.2 for feature counting and edgeR 3.26.0 for differential expression analysis.^66^ All scripts and data sets are or will be available at the following DOIs: 10.5281/zenodo.10694070 (data) and 10.5281/zenodo.10694150 (code).

## Acknowledgments

We thank Julien Luneau, Afra Salazar, Oliver Meacock, Massimo Amicone and Margaret Vogel for useful and constructive feedback on the manuscript. We thank Christopher van der Gast and Ian Thompson for the 4 species used. We thank Sabrina Rivera for help with the experiments. PP and SM conceived the study, PP, SM and MGG designed the experiment, PP performed the evolution experiment and all follow-up experiments with the help of SEAT, MGG and PP extracted the DNA and sent it for sequencing, EU performed bioinformatic analysis after some initial analysis by MGG, RDM performed RNA extractions, PP, EU and SM wrote the paper. PP and EU were funded by the University of Lausanne, MGG, RDM and SM were funded by European Research Council Starting Grant 715097, and SEAT and SM by the NCCR Microbiomes from the Swiss National Science Foundation.

## Supplementary tables and figures

**Figure S1:**
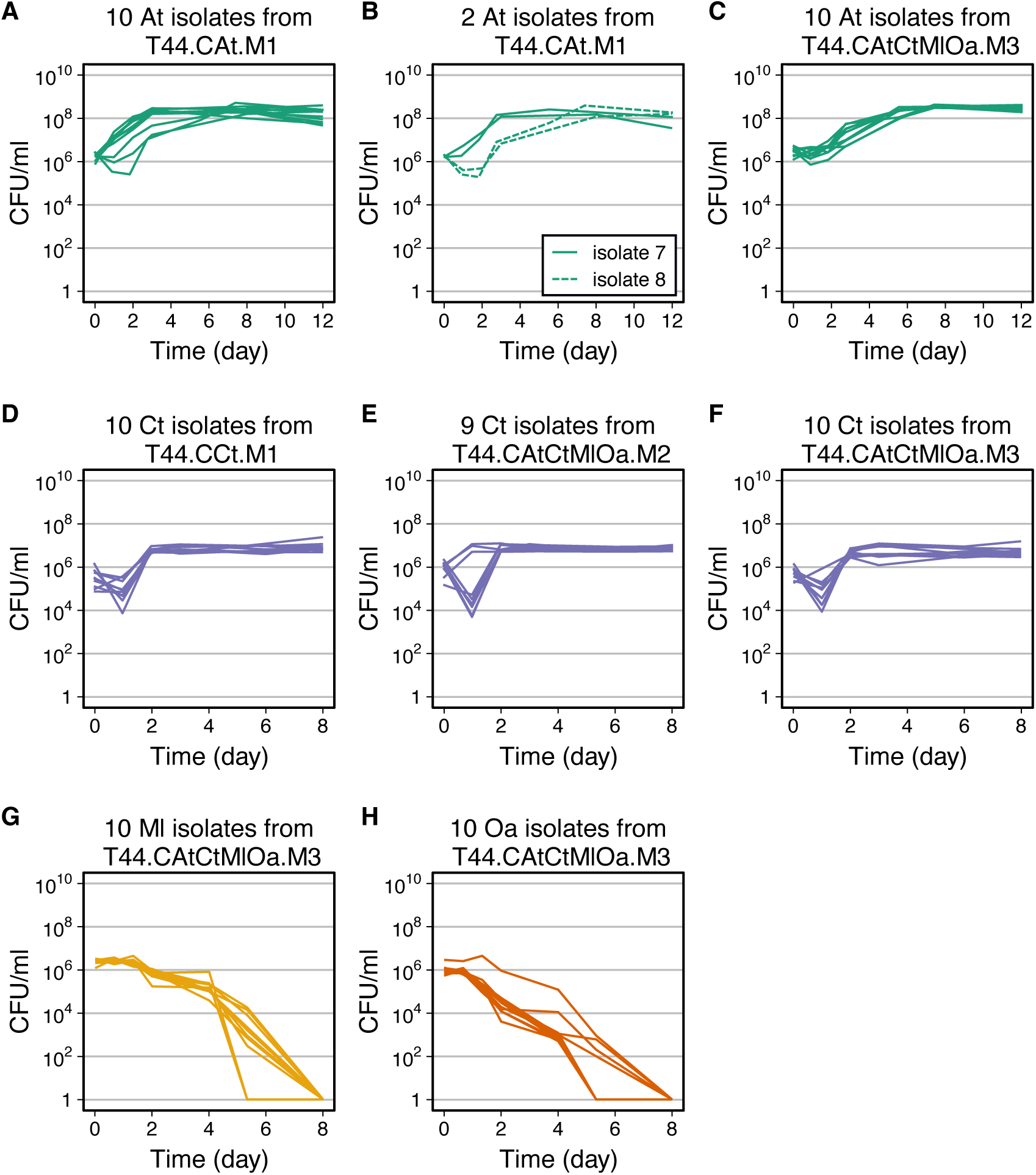
Growth curves of *A. tumefaciens* and *C. testosteroni* isolates from transfer 44. (A) Ten isolates of *A. tumefaciens* evolved alone from microcosm 1. (B) Two biological replicates of two of the isolates of *A. tumefaciens* shown in panel A to verify their growth differences. (C) Ten isolates of *A. tumefaciens* when evolved together with others (CAtCtMlOa, microcosm 3). (D) Ten isolates of *C. testosteroni* evolved alone from microcosm 1. (E) Nine isolates of *C. testosteroni* when evolved together with others (CAtCtMlOa, microcosm 2). This suggests some intra-species variability, which we investigate further in Fig. S2. (F) Ten isolates of *C. testosteroni* when evolved together with others (CAtCtMlOa, microcosm 3). (G) Ten isolates of *M. liquefaciens* when evolved together with others (CAtCtMlOa, microcosm 3). (H) Ten isolates of *O. anthropi* when evolved together with others (CAtCtMlOa, microcosm 3).

**Figure S2:**
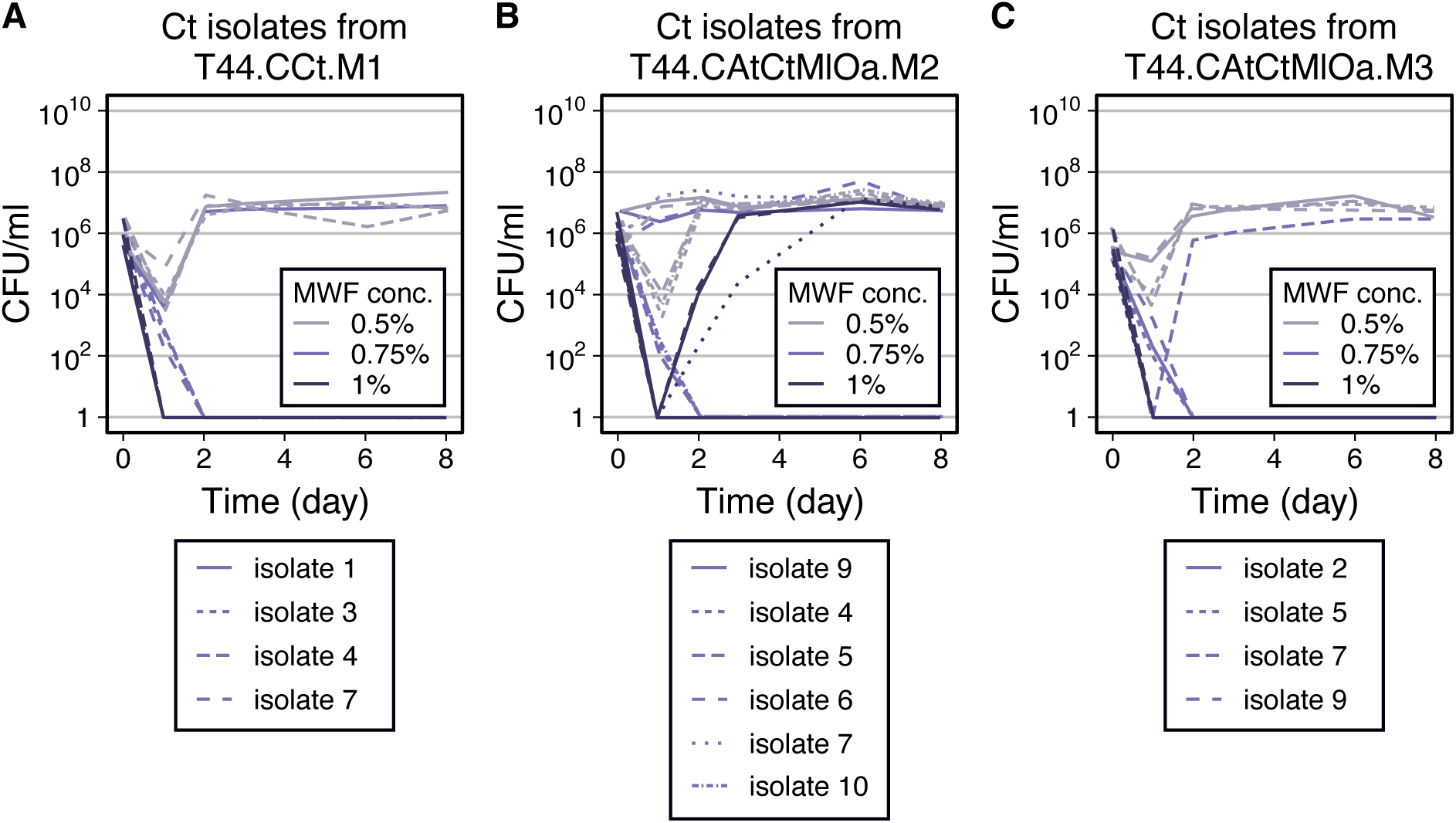
Growth curves of *C. testosteroni* isolates from transfer 44 in increasing concentrations of MWF over 8 days. All other experiments in this study were done at MWF concentration 0.5%. (A) Four isolates of *C. testosteroni* evolved alone, from microcosm 1. (B) Six isolates of *C. testosteroni* when evolved together with others (CAtCtMlOa) from microcosm 2. Here we see that some isolates are able to grow at higher MWF concentrations than we used in our experiment (0.5%) (C) Four isolates of *C. testosteroni* when evolved together with others (CAtCtMlOa) from microcosm 3.

**Figure S3:**
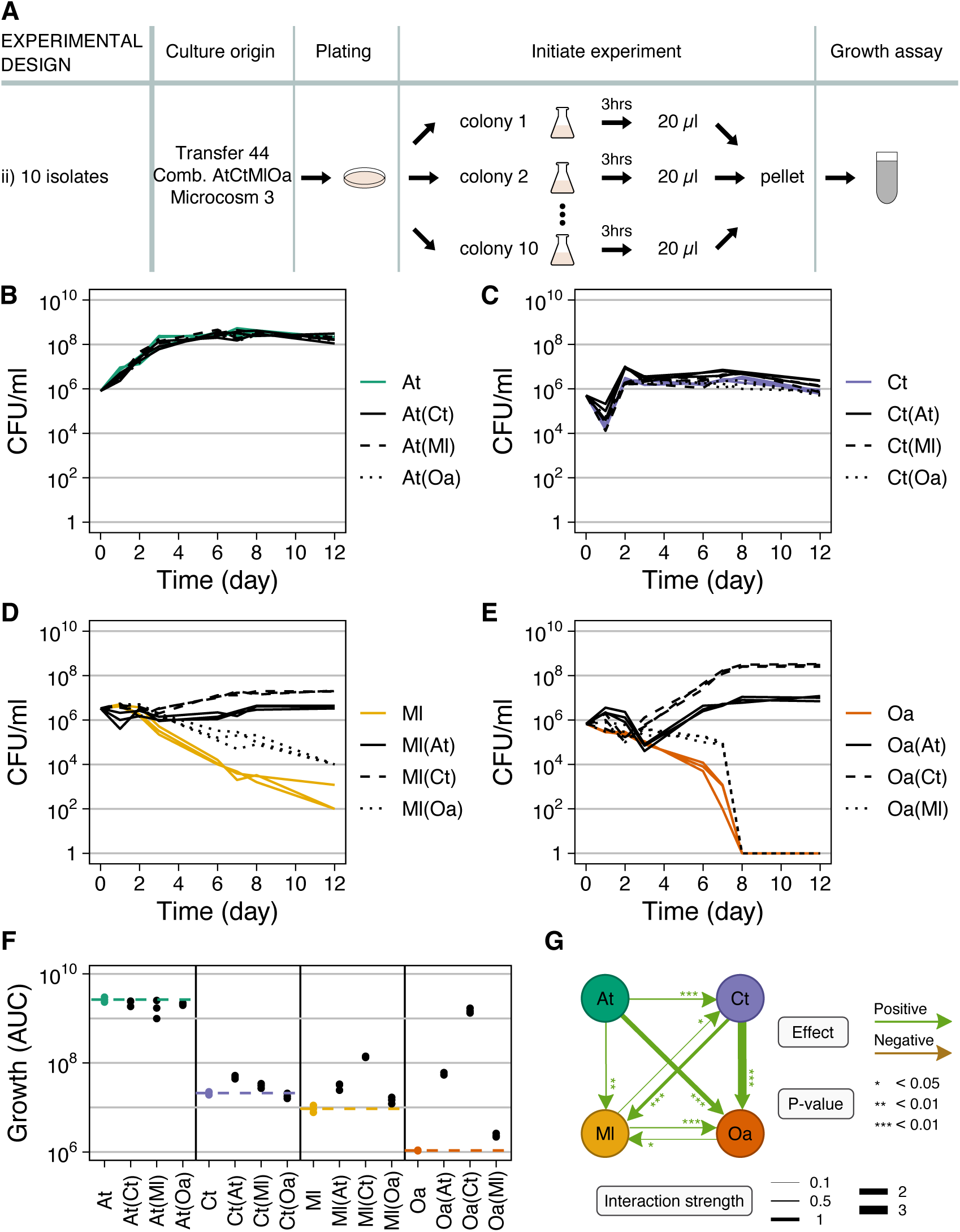
Comparison of co-evolved microcosm 3 mono- and pairwise co-cultures. (A) Ten evolved isolates of the same species were randomly picked and grown alone 3 hours to exponential phase, then washed, resuspended and mixed in equal proportions in MWF. (B=E) Population size quantified in colony-forming units per milliliter over time for mono-cultures (in color) and pairwise co-cultures (in black; co-culture partner indicated in brackets). In the co-cultures, each species could be quantified separately by selective plating. Each panel shows the data for 1 species: (B) *A. tumefaciens* (At), (C) *C. testosteroni* (Ct), (D) *M. liquefaciens* (Ml) and (E) *O. anthropi* (Oa). (F) AUC in B=E. Dashed lines indicate the mean of the mono-cultures, shown in color. Statistical significance and interaction strengths data are shown in Dataset S1.

**Figure S4:**
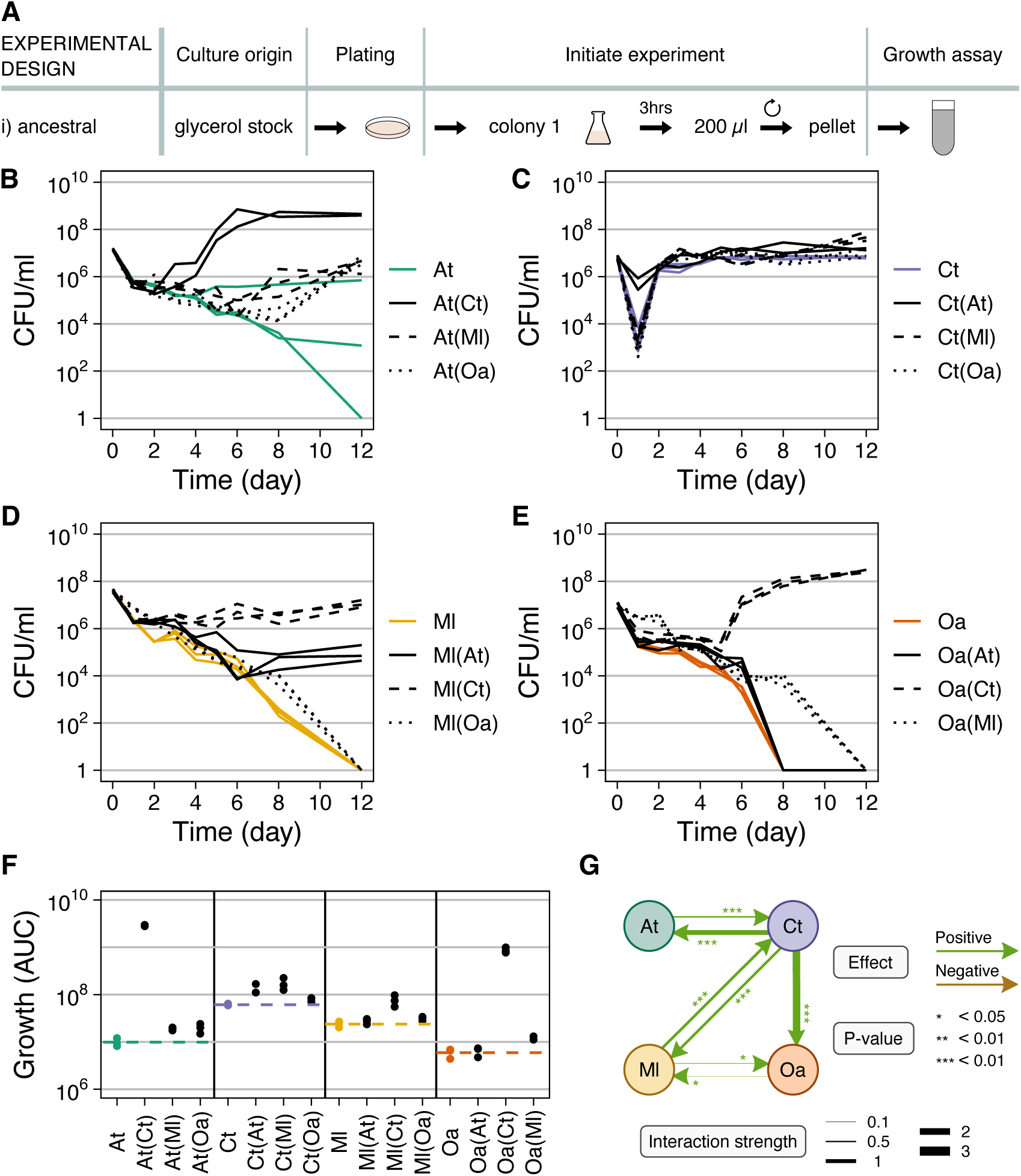
Comparison of ancestral mono- and pairwise co-cultures, adapted from ^34^. (A) Glycerol stock of ancestral isolate was grown alone 3 hours to exponential phase, then washed and resuspended in MWF. (B=E) Population size quantified in colony-forming units per milliliter over time for mono-cultures (in color) and pairwise co-cultures (in black; co-culture partner indicated in brackets). In the cocultures, each species could be quantified separately by selective plating. Each panel shows the data for 1 species: (B) *A. tumefaciens* (At), (C) *C. testosteroni* (Ct), (D) *M. liquefaciens* (Ml) and (E) *O. anthropi* (Oa). (F) AUC in B=E. Dashed lines indicate the mean of the mono-cultures, shown in color.

**Figure S5:**
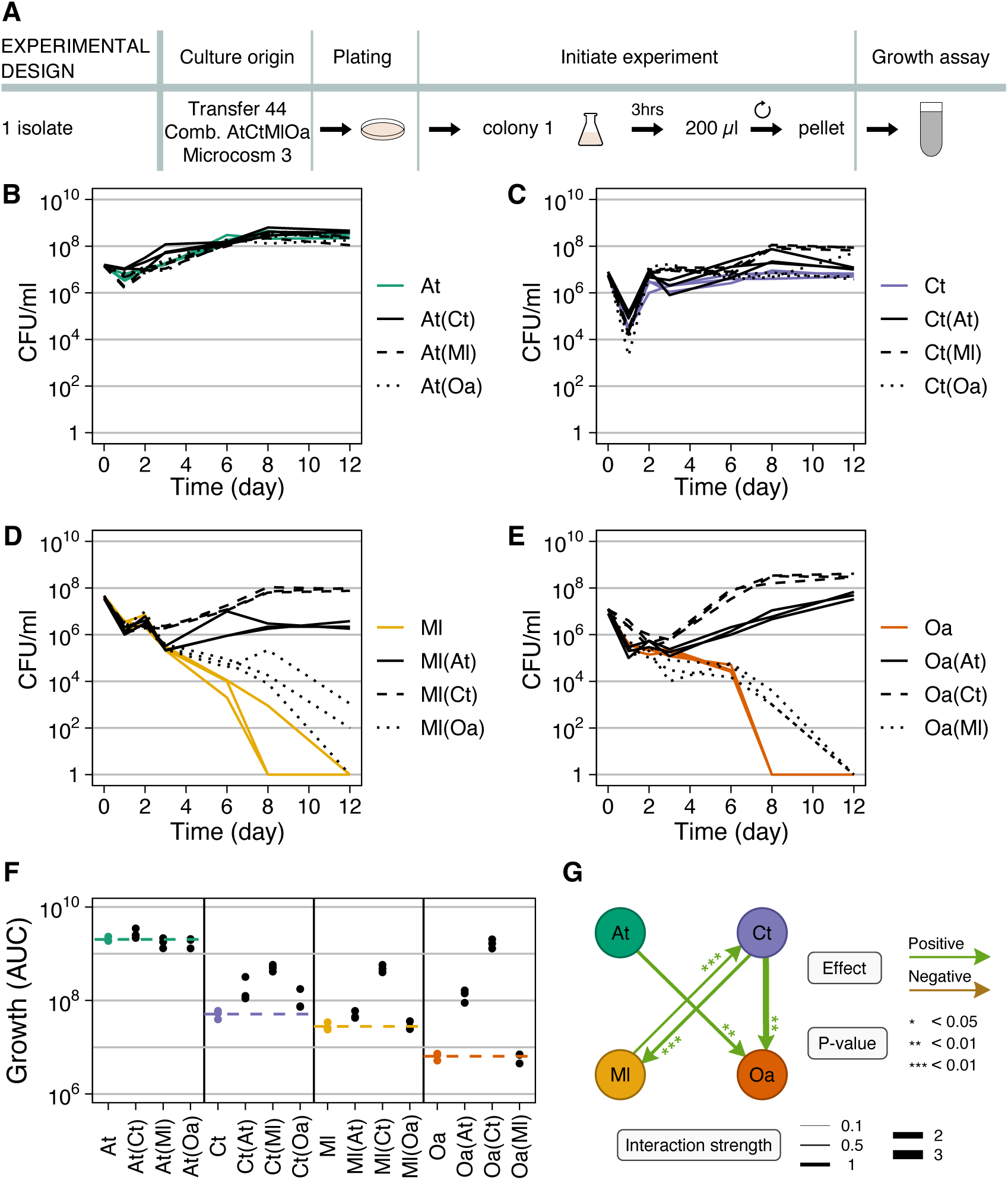
Comparison of co-evolved microcosm 3 mono- and pairwise co-cultures. (A) One evolved isolate of each species was randomly picked and grown alone 3 hours to exponential phase, then washed, resuspended and mixed in equal proportions in MWF. (B=E) Population size quantified in colony-forming units per milliliter over time for mono-cultures (in color) and pairwise co-cultures (in black; co-culture partner indicated in brackets). In the co-cultures, each species could be quantified separately by selective plating. Each panel shows the data for 1 species: (B) *A. tumefaciens* (At), (C) *C. testosteroni* (Ct), (D) *M. liquefaciens* (Ml) and (E) *O. anthropi* (Oa). (F) AUC in B=E. Dashed lines indicate the mean of the mono-cultures, shown in color. Statistical significance and interaction strengths data are shown in Dataset S2.

**Figure S6:**
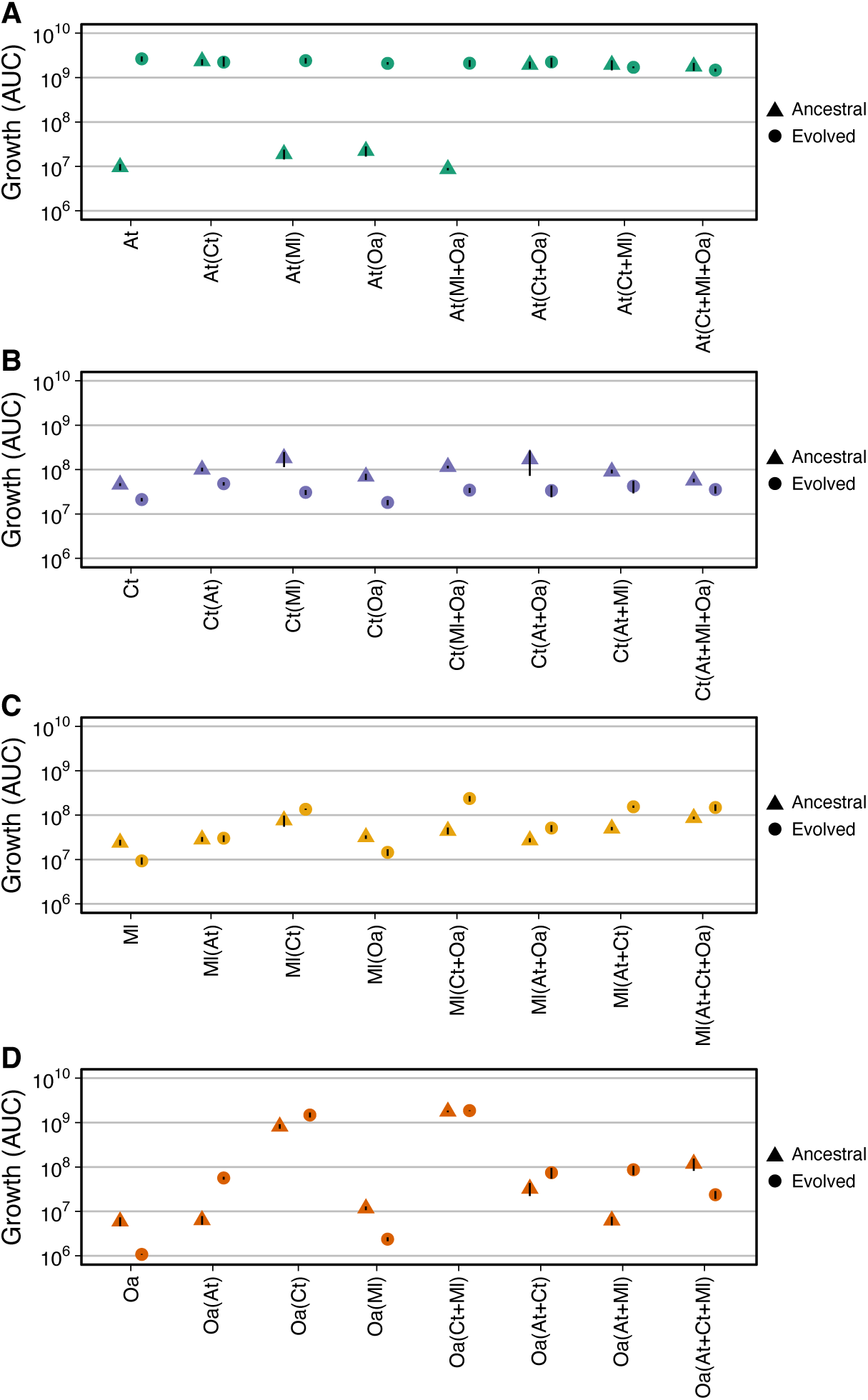
AUC comparison of ancestral species and those evolved in CAtCtMlOa, microcosm 3, including mono- and co-cultures treatments for (A) *A. tumefaciens*, (B) *C. testosteroni*, (C) *M. liquefaciens*, and (D) *O. anthropi*. Evoled strains were co-culured with isolates from the same microcosm and ancestral strains were co-cultured with other ancestors.

**Figure S7:**
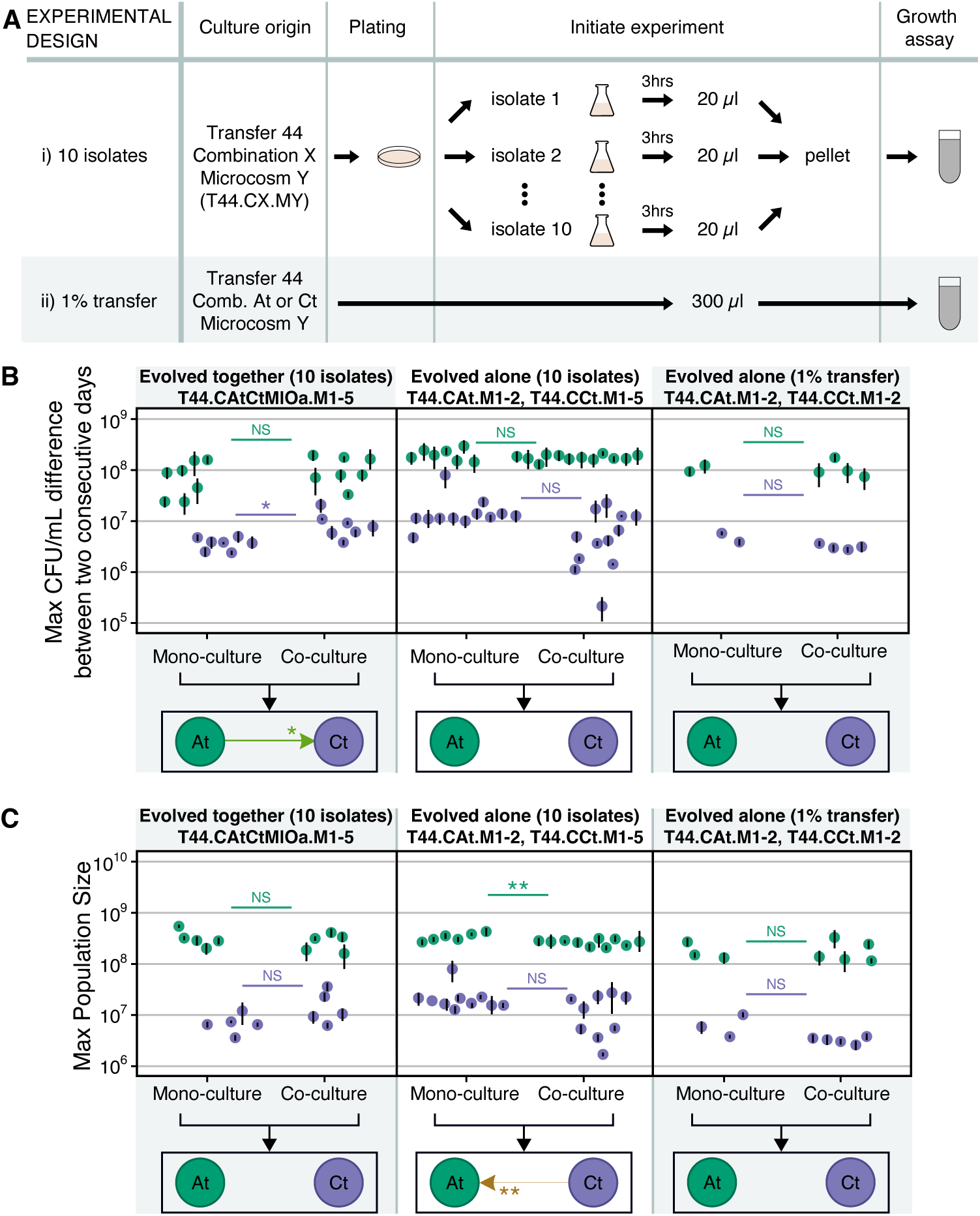
Interactions based on maximum growth rate and maximum population size. A) Protocols for growth assays, matching those in Fig. 3A. (B-C) Interactions between *A. tumefaciens* and *C. testosteroni* based on maximum growth rate quantified as the maximal CFU/ml difference between two consecutive days (B) or maximum population size (C), either evolved together (first column, CAtCtMlOa) or evolved alone (2nd and 3rd column, CAt and CCt, protocols i and ii from panel A) during 8-day growth assays. Other details are as in Fig. 3C.

**Figure S8:**
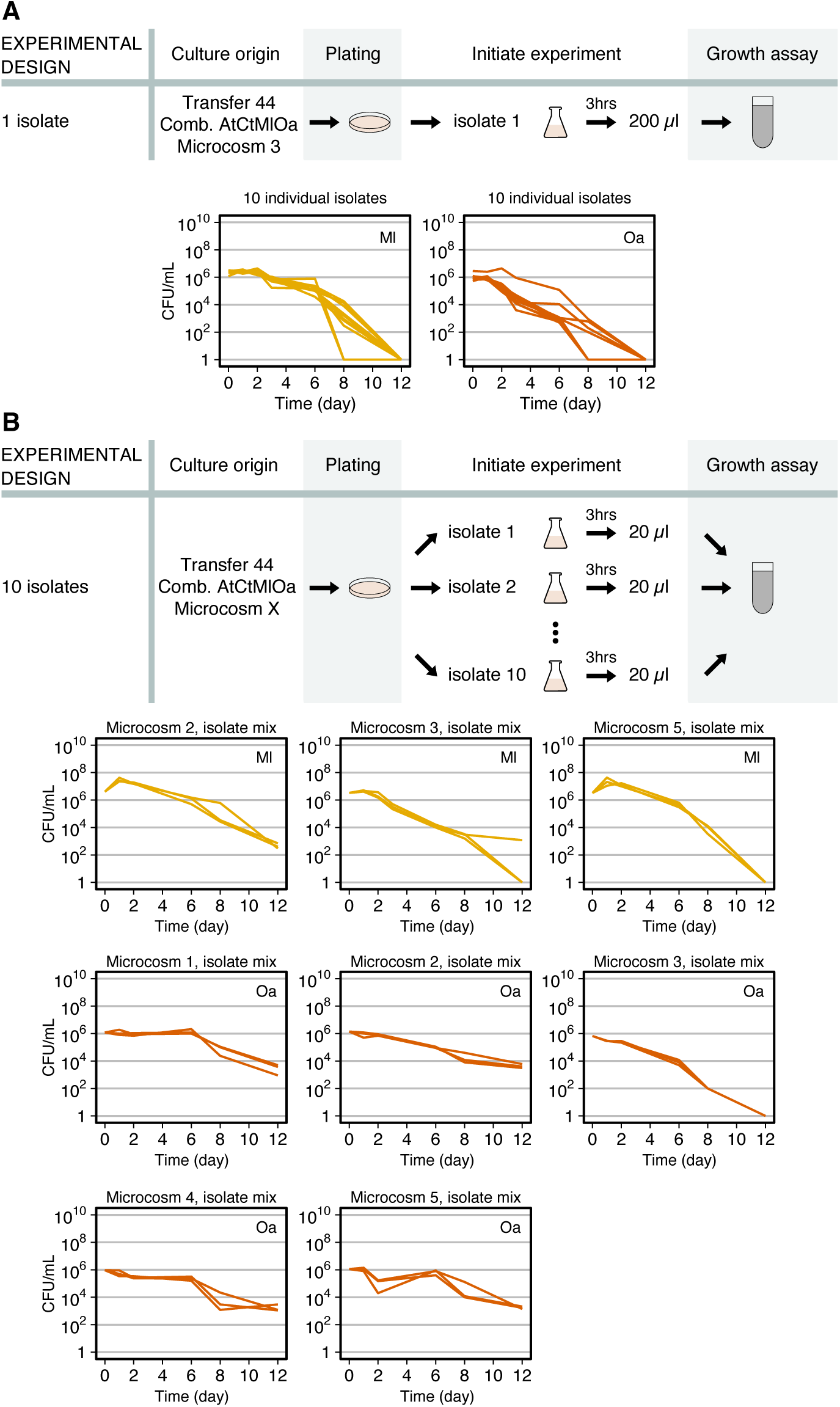
Mono-culture growth curves of evolved *M. liquefaciens* or *O anthropi* from transfer 44, CAtCtMlOa during 12-day growth assays. Conditions and microcosms are indicated above each graph. (A) One isolates was randomly picked and grown alone 3 hours to exponential phase, then washed and resuspended in MWF. Each growth curve represents one of 10 such isolates. (B) Ten evolved isolates were randomly picked and grown alone 3 hours to exponential phase, then washed, resuspended as a mixed culture in MWF. Each panel shows triplicates of isolates the same condition and microcosm.

**Figure S9:**
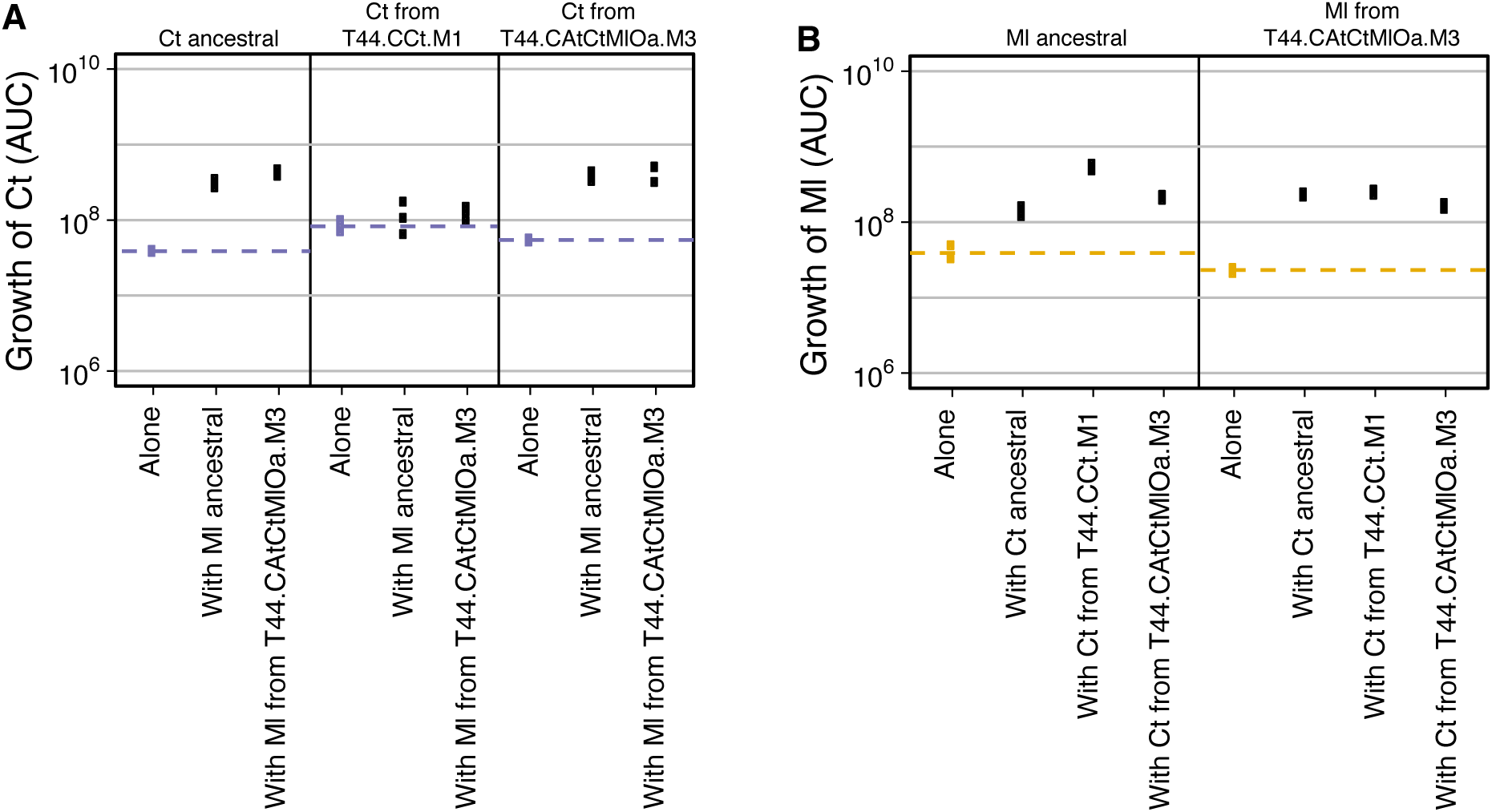
Interactions between *Ct* and *Ml*. (A) Growth of different *Ct* isolates (ancestral, evolved alone or evolved with the three others) alone or in co-culture with different *Ml* isolates (ancestral or evolved with the three others). (B) Growth of different *Ml* isolates alone or in co-culture with different *Ct* isolates. Community-evolved *Ct* and *Ml* were isolated from the same microcosm. Ancestral and community-evolved *Ct* and *Ml* all had positive effects on one another, but the positive effects did not increase between the isolates of the two species coming from the same microcosm, suggesting that at least in this microcosm, *Ct* and *Ml* did not evolve stronger mutualism.

**Figure S10:**
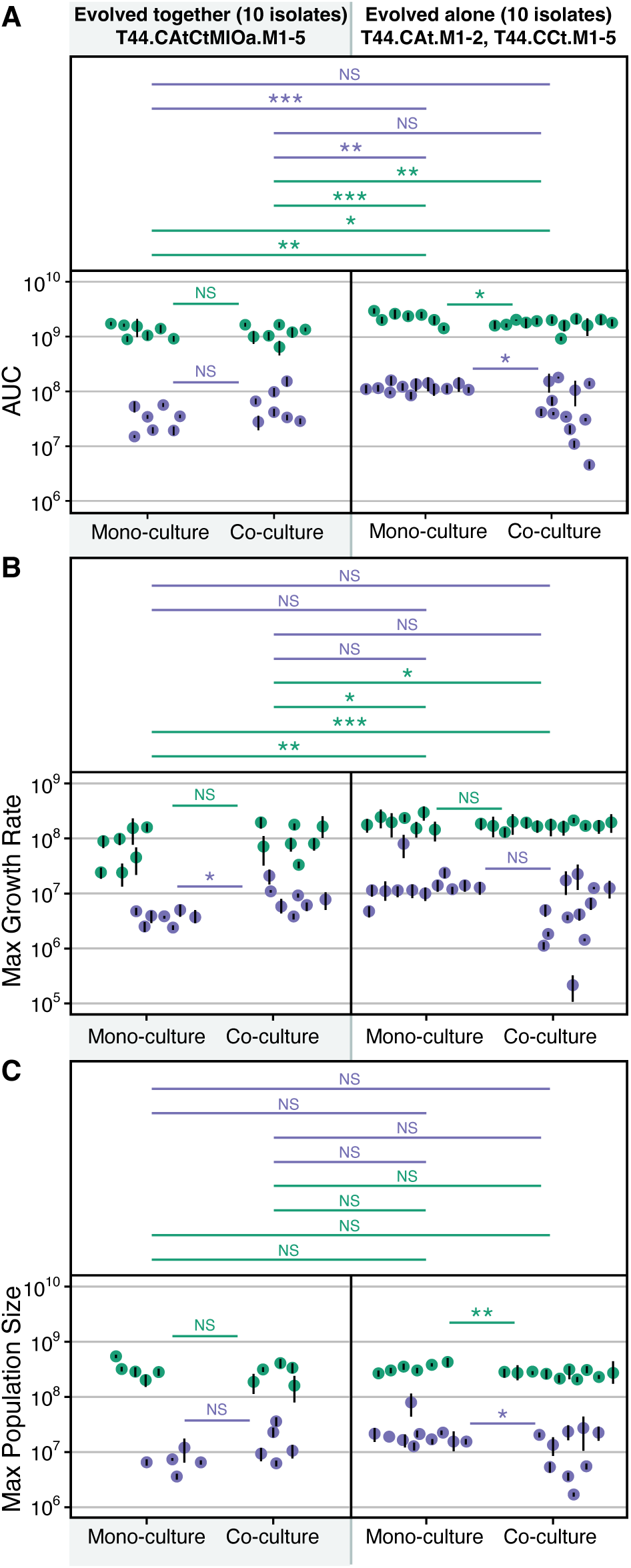
Inter-group comparison from Fig. 3C. The data show interactions between *A. tumefaciens* and *C. testosteroni* co-evolved (first column) or mono-evolved (second column) during 8-day growth assays. The first row measures the AUC of their growth curves during 8-day growth assays. The second and third row measure their maximum growth rates and maximum population size reached during these growth assays.

**Figure S11:**
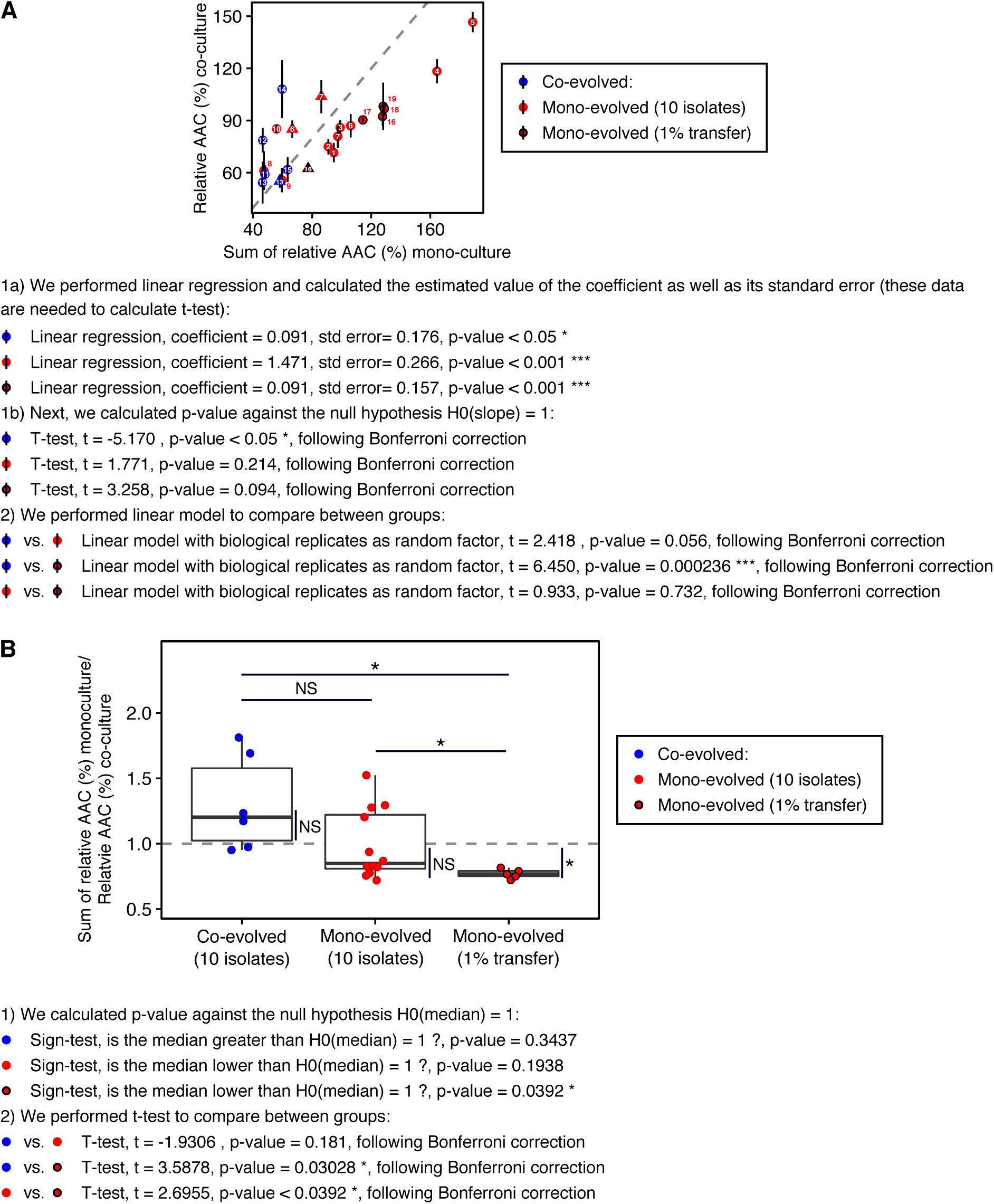
Results of statistical analysis of the additive null model to degradation efficiency in Fig. 5C). (A) Linear model. (B) T-test. Co-evolved is from species combination CAtCtMlOa and mono-evolved from CAt or CCt.

**Figure S12:**
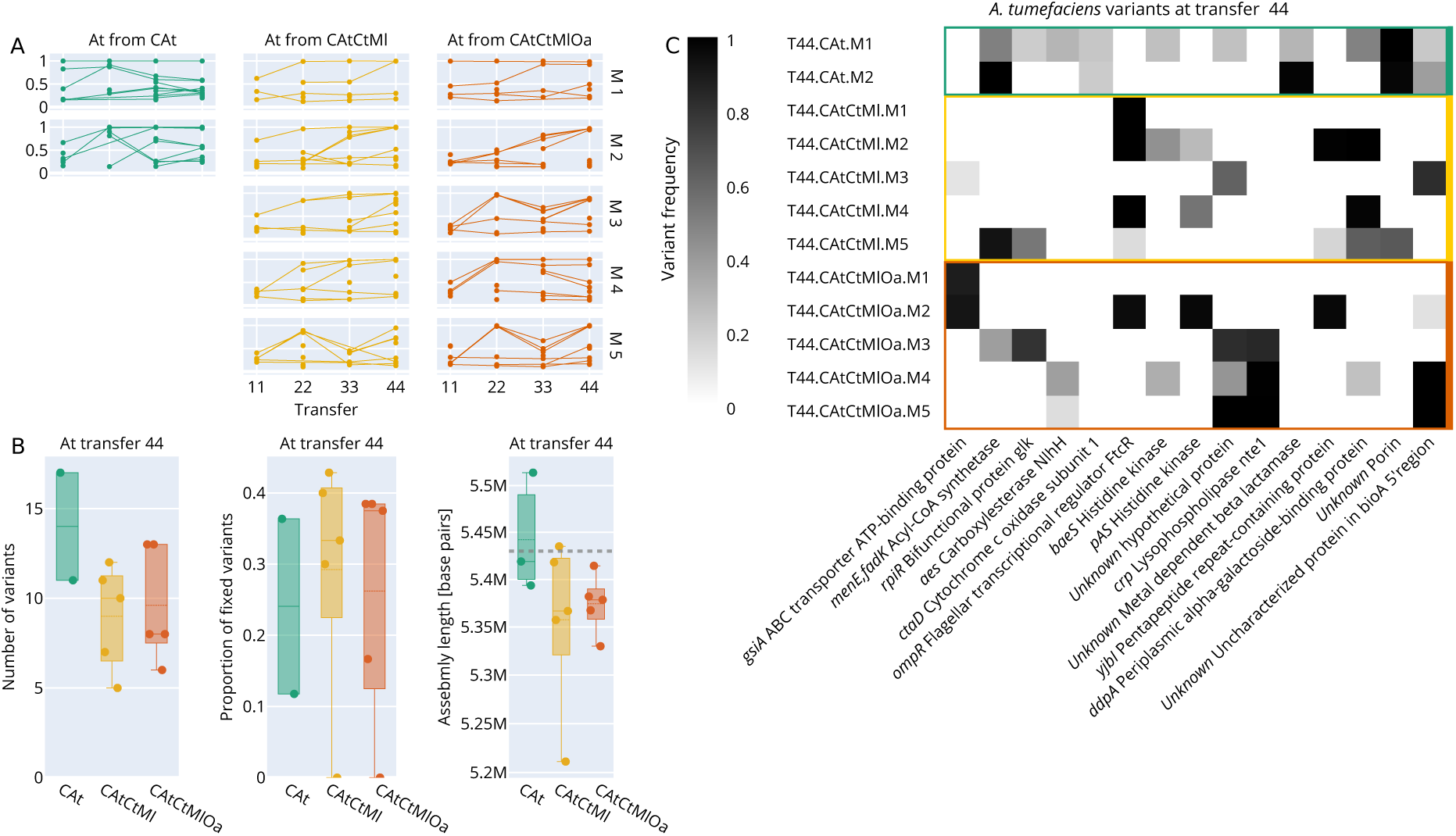
(A) Variant frequency trajectories in all *A. tumefaciens* populations. (B) Number of variants found in each *A. tumefaciens* population (left). De-novo long-read assembly lengths of selected isolates. Dashed line represents assembly length of the ancestor (middle). Proportion of variants that reached fixation (right). (C) Mutated genes with protein annotation that were found in at least two *A. tumefaciens* populations. The color indicates the frequency of the mutated allele.

**Figure S13:**
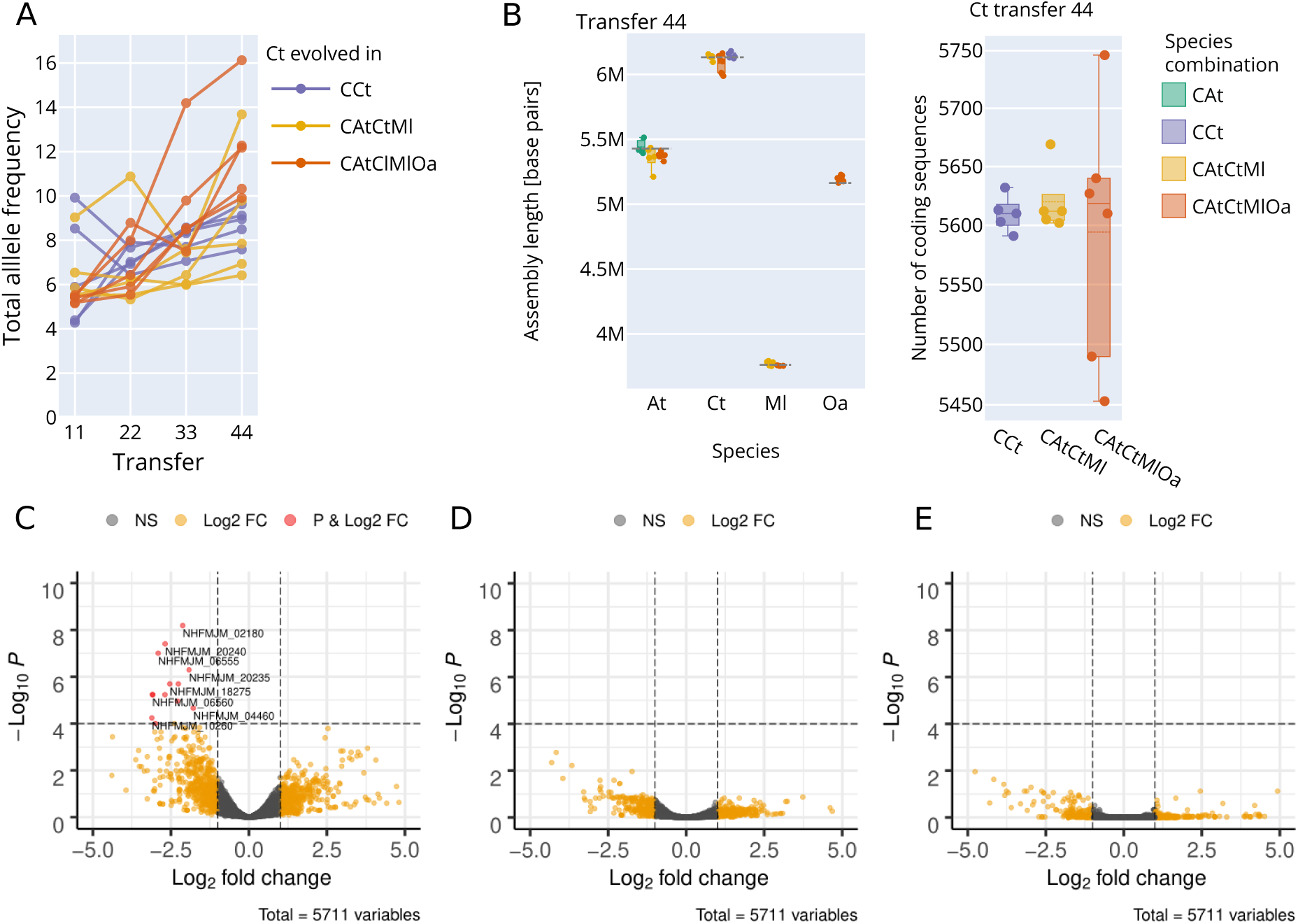
(A) Total allele frequency for *C. testosteroni*. (B) Long-read assembly lengths of isolates from transfer 44 with the dashed line representing the assembly length of the ancestor (left). Number of coding sequences per assembly (right). (C) Gene expression for *C. testosteroni* evolved under condtition 2 compared to ancestor. (D) Gene expression for *C. testosteroni* evolved under condtion 3 compared to ancestor. (E) Gene expression for *C. testosteroni* evolved under CAtCtMlOa compared to ancestor.

**Figure S14:**
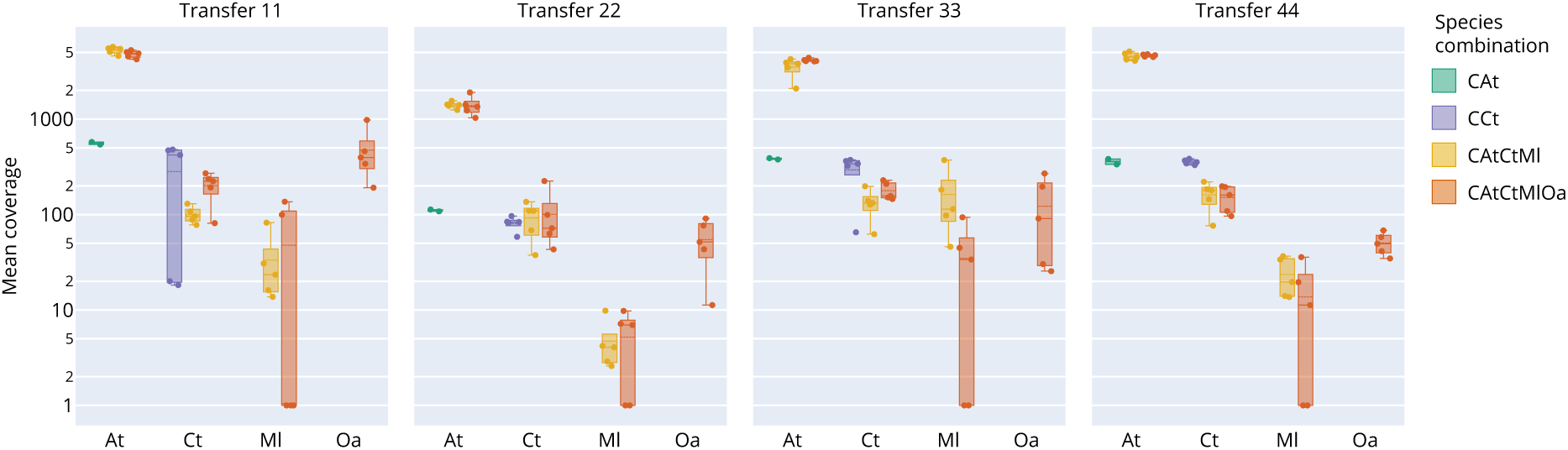
Mean Illumina coverage.

**Figure S15:**
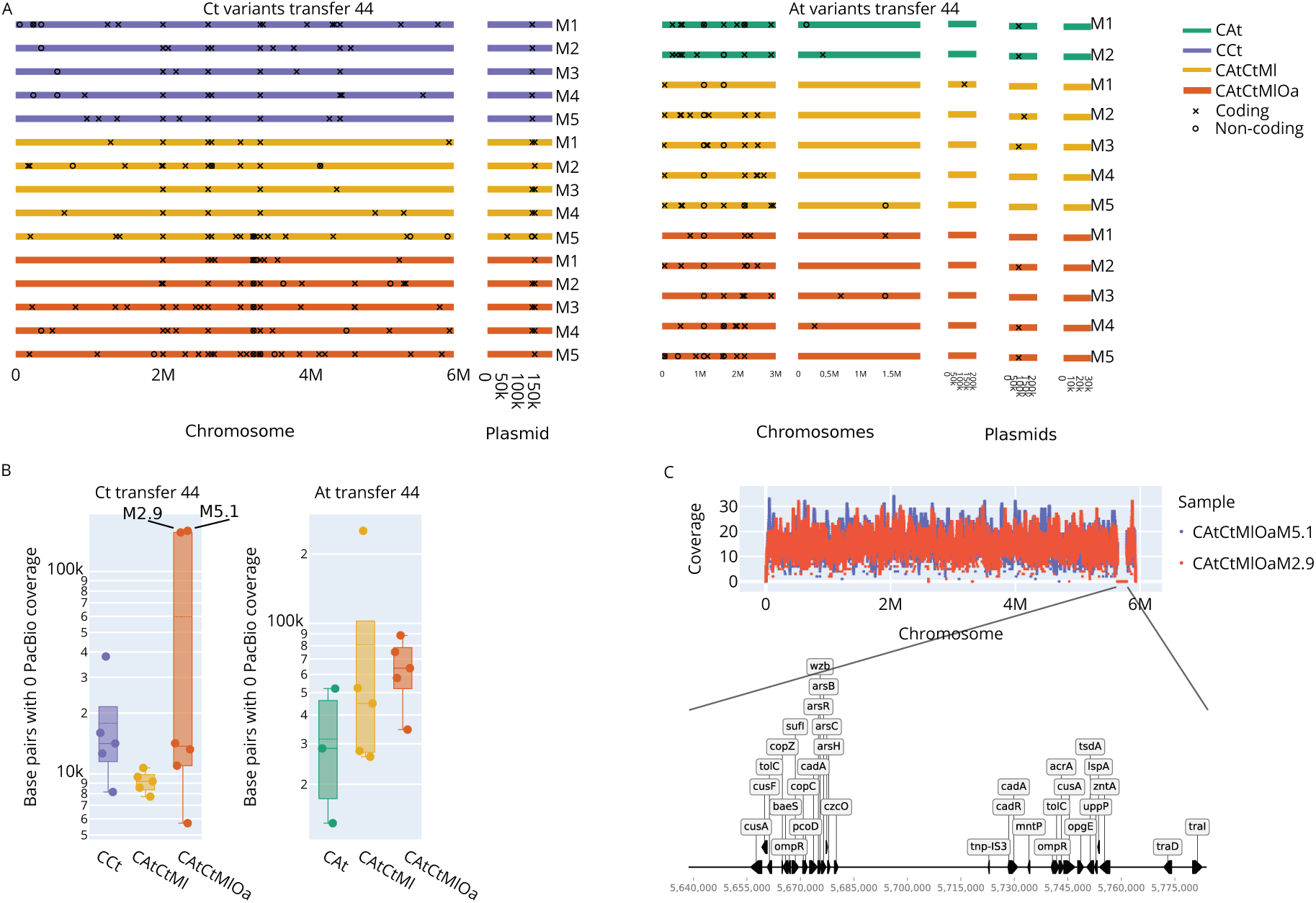
(A) Positions of variants across all frequencies identified from the Illumina data from the last transfer. (B) Base pairs with zero-coverage when aligning corrected PacBio reads to the reference genome. (C) PacBio coverage for two Ct isolates of CAtCtMlOa showing large deletion (top). Annotation of deleted sequence (bottom).

**Figure S16:**
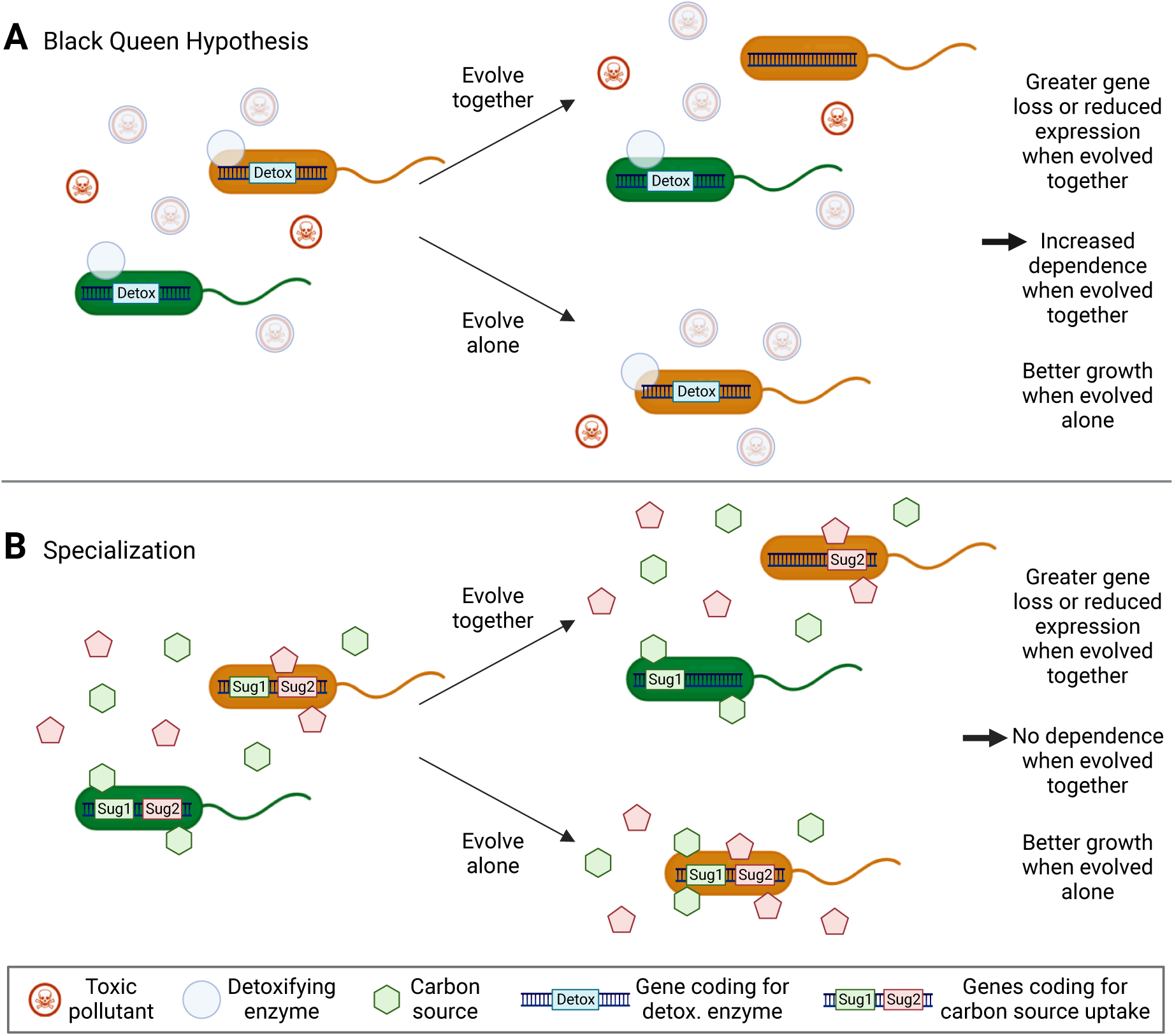
BQH and specialization make similar predictions. (A) The BQH predicts that species evolved together in community should lose traits coding for public goods, like detoxification genes either by deletions or mutations leading to reduced gene expression. Such losses should not be observed when evolving alone. Species evolved together should therefore grow significantly worse alone and depend on the partner species for survival. (B) The evolution of specialization predicts similar trait loss when evolving together and should similarly grow best when evolved alone, but species evolved in community should not depend on their partners to grow alone (black arrows on the right). Initially both species can take up both carbon sources but with a preference for one or the other. After evolution alone, the orange species can take up both efficiently. Generated using Biorender.

**Table S1:**
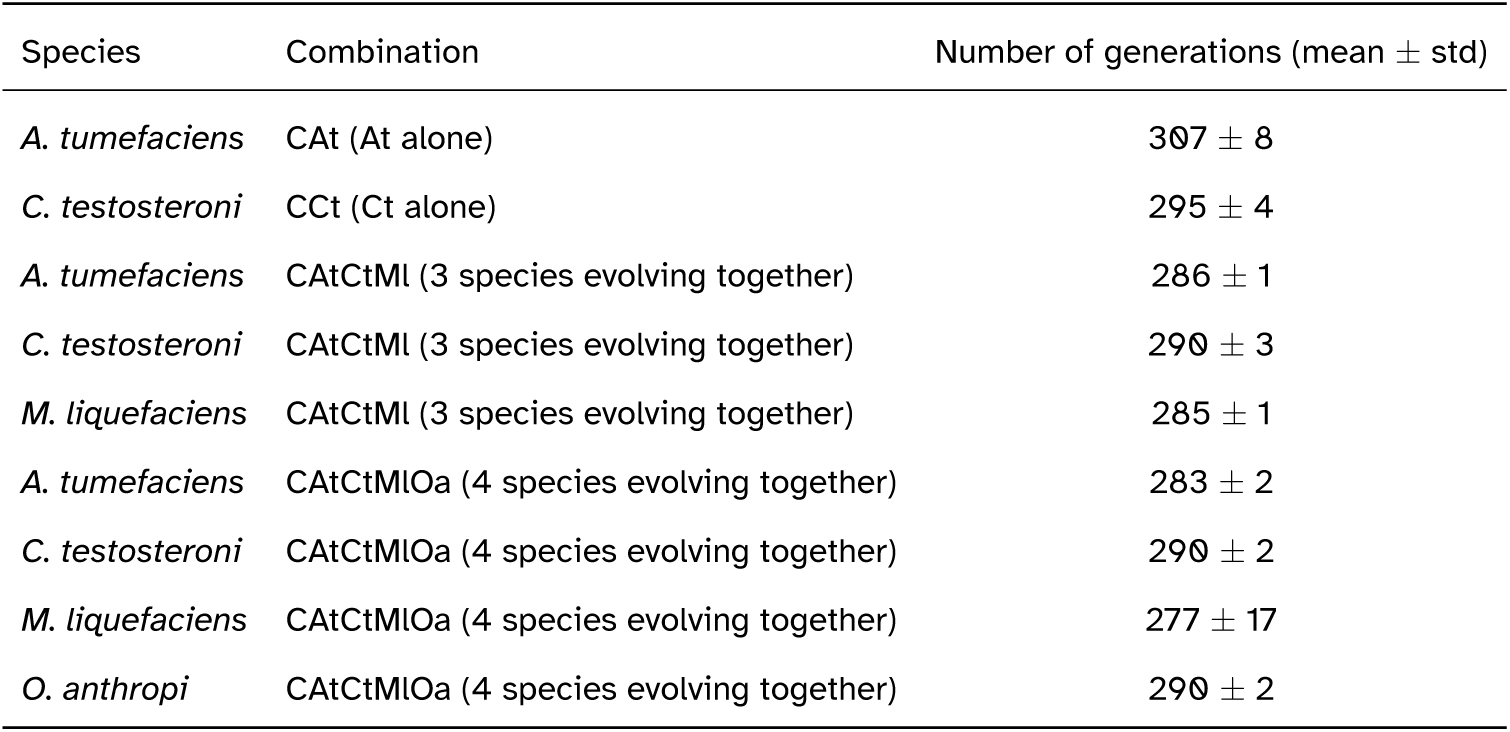
Number of generations per species averaged over microcosms in which that species survived until transfer 44. The number of generations *n* was computed for each microcosm and each transfer as *n* = *log*_10_(*b*/*B*)/*log*_10_(2), where *b* is the CFU/ml at the beginning of a transfer (CFU/ml of the previous transfer divided by 100) and *B* the CFU/ml at the end of that same transfer. We then summed *n* over all transfers and took the average over all microcosms of that species in a given combination.

| Species | Combination | Number of generations (mean ± std) |
| --- | --- | --- |
| *A. tumefaciens* | CAt (At alone) | 307 ± 8 |
| *C. testosteroni* | CCt (Ct alone) | 295 ± 4 |
| *A. tumefaciens* | CAtCtMl (3 species evolving together) | 286 ± 1 |
| *C. testosteroni* | CAtCtMl (3 species evolving together) | 290 ± 3 |
| *M. liquefaciens* | CAtCtMl (3 species evolving together) | 285 ± 1 |
| *A. tumefaciens* | CAtCtMlOa (4 species evolving together) | 283 ± 2 |
| *C. testosteroni* | CAtCtMlOa (4 species evolving together) | 290 ± 2 |
| *M. liquefaciens* | CAtCtMlOa (4 species evolving together) | 277 ± 17 |
| *O. anthropi* | CAtCtMlOa (4 species evolving together) | 290 ± 2 |

**Table S2:**
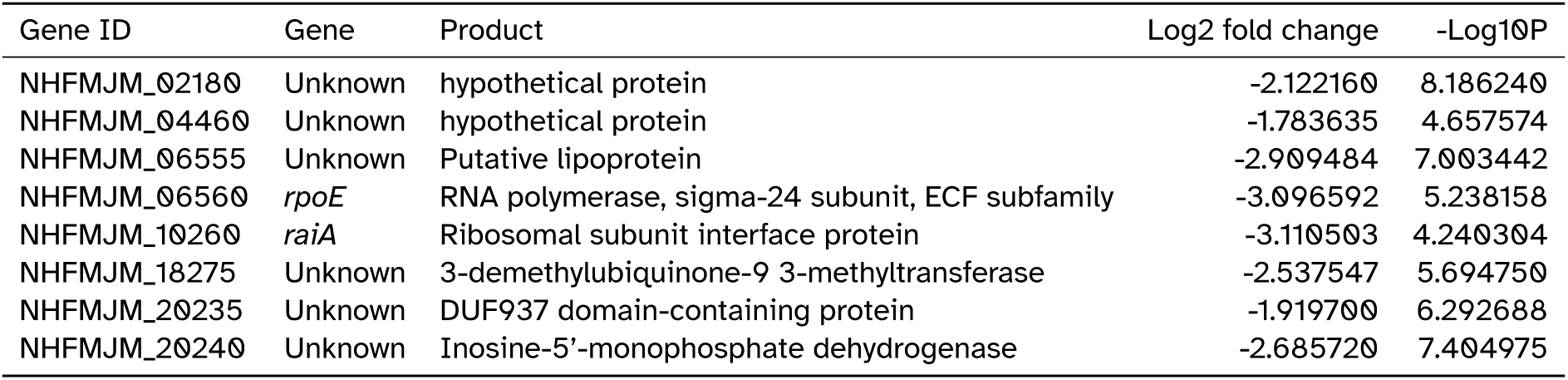
Differentially expressed genes in *C. testosteroni* evolved alone (CCt).

## Notes

https://zenodo.org/uploads/10694070

https://zenodo.org/records/10694151

### Competing Interest Statement

The authors have declared no competing interest.

### Summary of Updates

A few changes were made to the abstract (e.g. full species name written out) and one header in the results section.

