## Supplementary figures and tables for "The evolution of reduced facilitation in a four-species bacterial community"

### Supplementary tables and figures

| Species | Combination | Number of generations (mean $\pm$ std) |
|---|---|---|
| *A. tumefaciens* | CAt (At alone) | $307 \pm 8$ |
| *C. testosteroni* | CCt (Ct alone) | $295 \pm 4$ |
| *A. tumefaciens* | CAtCtMl (3 species evolving together) | $286 \pm 1$ |
| *C. testosteroni* | CAtCtMl (3 species evolving together) | $290 \pm 3$ |
| *M. liquefaciens* | CAtCtMl (3 species evolving together) | $285 \pm 1$ |
| *A. tumefaciens* | CAtCtMlOa (4 species evolving together) | $283 \pm 2$ |
| *C. testosteroni* | CAtCtMlOa (4 species evolving together) | $290 \pm 2$ |
| *M. liquefaciens* | CAtCtMlOa (4 species evolving together) | $277 \pm 17$ |
| *O. anthropi* | CAtCtMlOa (4 species evolving together) | $290 \pm 2$ |

Table S1: Number of generations per species averaged over microcosms in which that species survived until transfer 44. The number of generations $n$ was computed for each microcosm and each transfer as $n = log_{10}(b/B)/log_{10}(2)$, where $b$ is the CFU/ml at the beginning of a transfer (CFU/ml of the previous transfer divided by $100$) and $B$ the CFU/ml at the end of that same transfer. We then summed $n$ over all transfers and took the average over all microcosms of that species in a given combination.

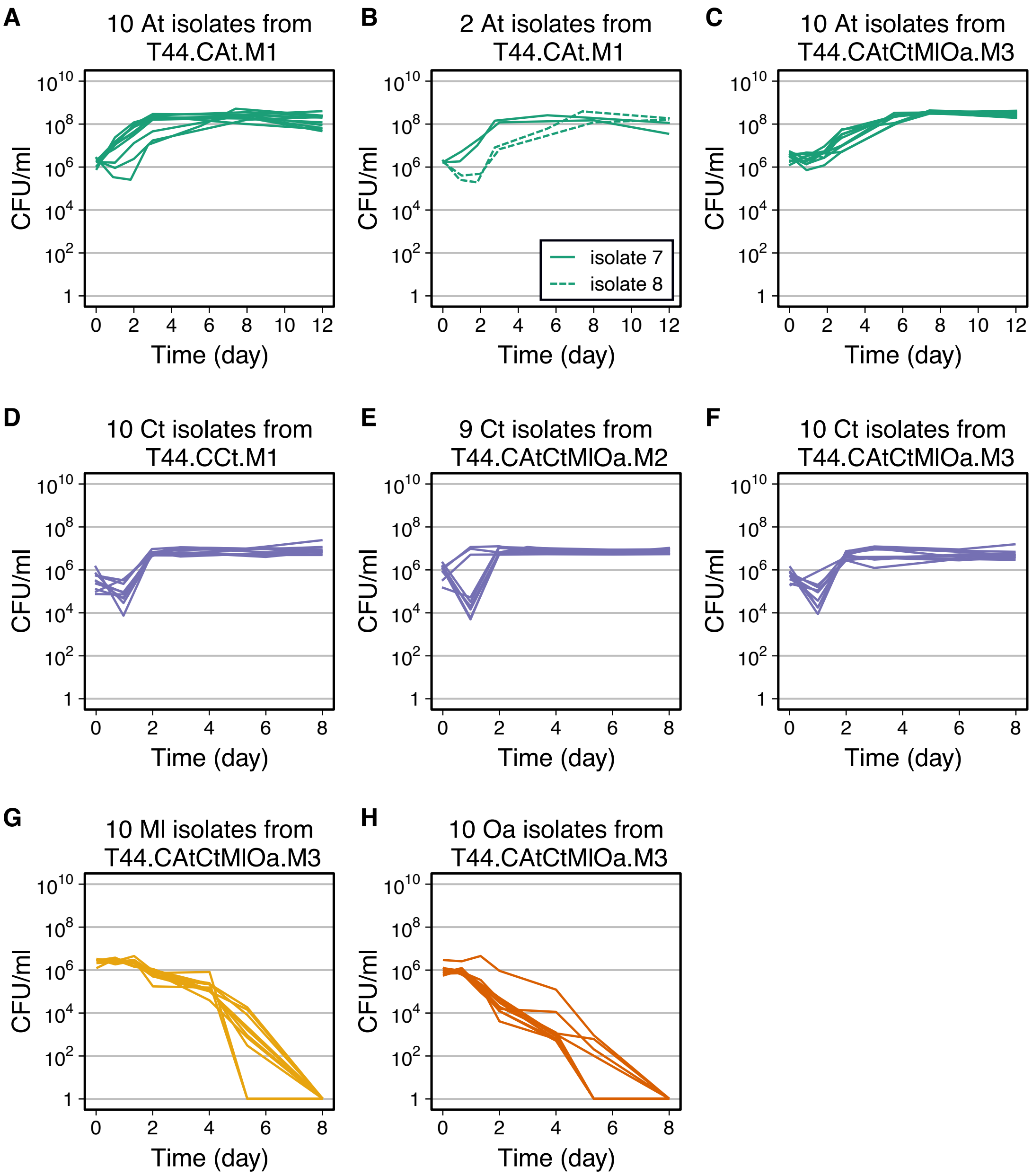


Figure S1: Growth curves of *A. tumefaciens* and *C. testosteroni* isolates from transfer 44. (A) Ten isolates of *A. tumefaciens* evolved alone from microcosm 1. (B) Two biological replicates of two of the isolates of *A. tumefaciens* shown in panel A to verify their growth differences. (C) Ten isolates of *A. tumefaciens* when evolved together with others (CAtCtMlOa, microcosm 3). (D) Ten isolates of *C. testosteroni* evolved alone from microcosm 1. (E) Nine isolates of *C. testosteroni* when evolved together with others (CAtCtMlOa, microcosm 2). This suggests some intra-species variability, which we investigate further in Fig. S2. (F) Ten isolates of *C. testosteroni* when evolved together with others (CAtCtMlOa, microcosm 3). (G) Ten isolates of *M. liquefaciens* when evolved together with others (CAtCtMlOa, microcosm 3). (H) Ten isolates of *O. anthropi* when evolved together with others (CAtCtMlOa, microcosm 3).

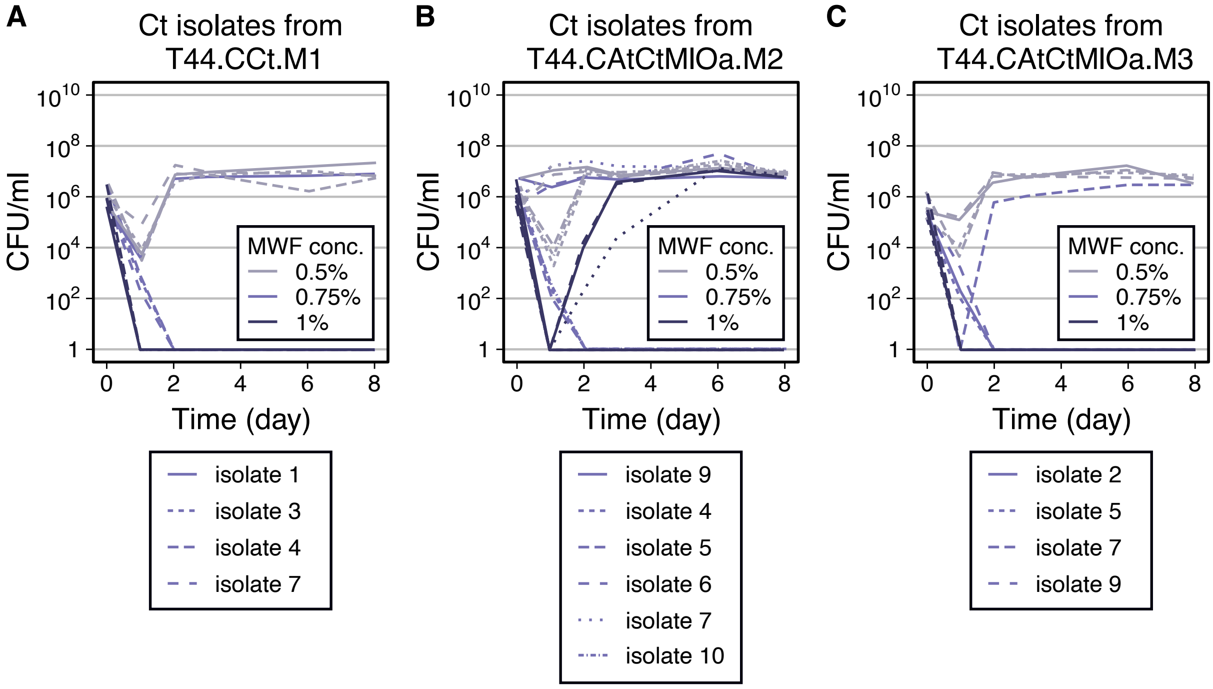


Figure S2: Growth curves of *C. testosteroni* isolates from transfer 44 in increasing concentrations of MWF over 8 days. All other experiments in this study were done at MWF concentration 0.5%. (A) Four isolates of *C. testosteroni* evolved alone, from microcosm 1. (B) Six isolates of *C. testosteroni* when evolved together with others (CAtCtMlOa) from microcosm 2. Here we see that some isolates are able to grow at higher MWF concentrations than we used in our experiment (0.5%) (C) Four isolates of *C. testosteroni* when evolved together with others (CAtCtMlOa) from microcosm 3.

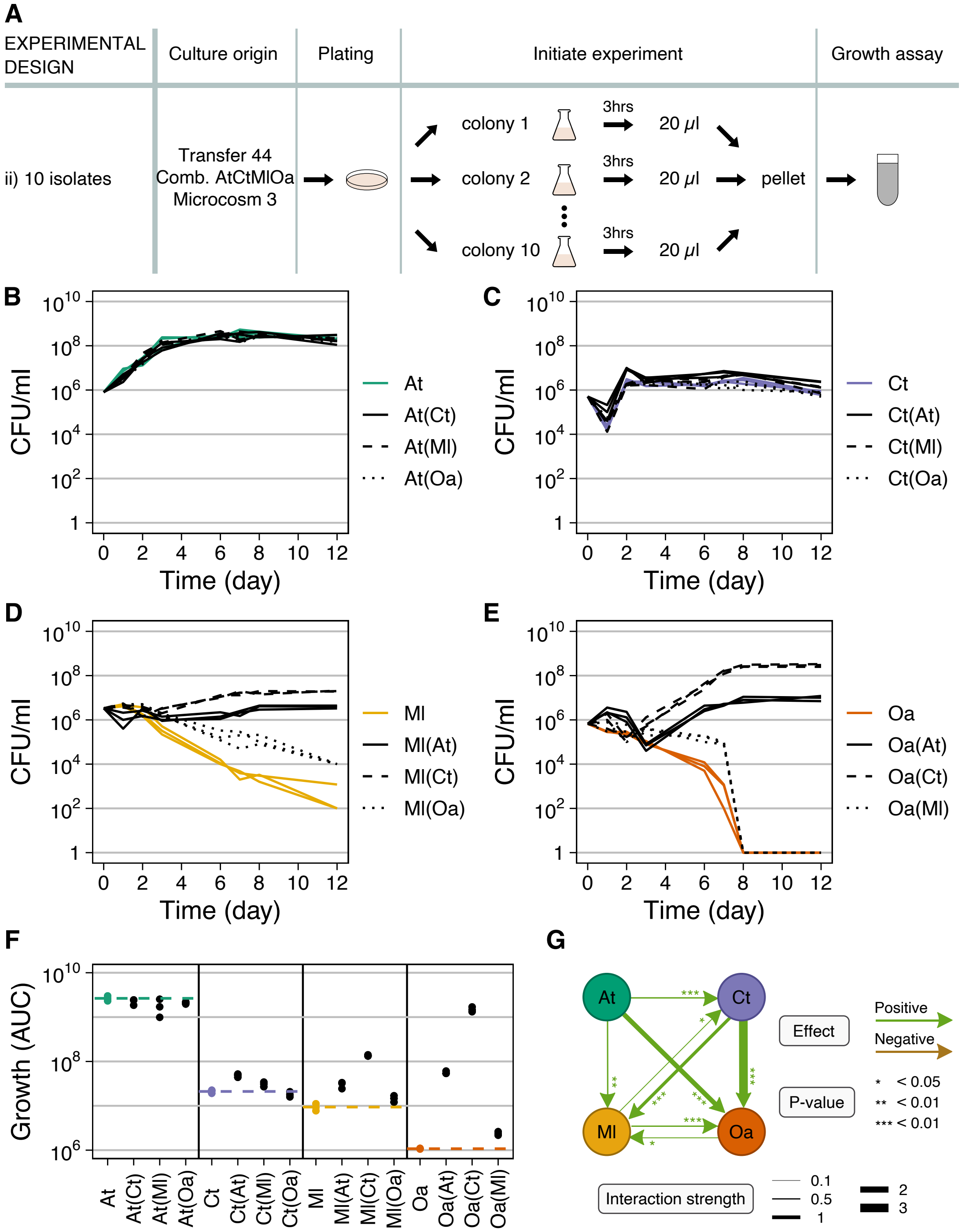


Figure S3: Comparison of co-evolved microcosm 3 mono- and pairwise co-cultures. (A) Ten evolved isolates of the same species were randomly picked and grown alone 3 hours to exponential phase, then washed, resuspended and mixed in equal proportions in MWF. (B=E) Population size quantified in colony-forming units per milliliter over time for mono-cultures (in color) and pairwise co-cultures (in black; co-culture partner indicated in brackets). In the co-cultures, each species could be quantified separately by selective plating. Each panel shows the data for 1 species: (B) *A. tumefaciens* (At), (C) *C. testosteroni* (Ct), (D) *M. liquefaciens* (Ml) and (E) *O. anthropi* (Oa). (F) AUC in B=E. Dashed lines indicate the mean of the mono-cultures, shown in color. Statistical significance and interaction strengths data are shown in Dataset S1.

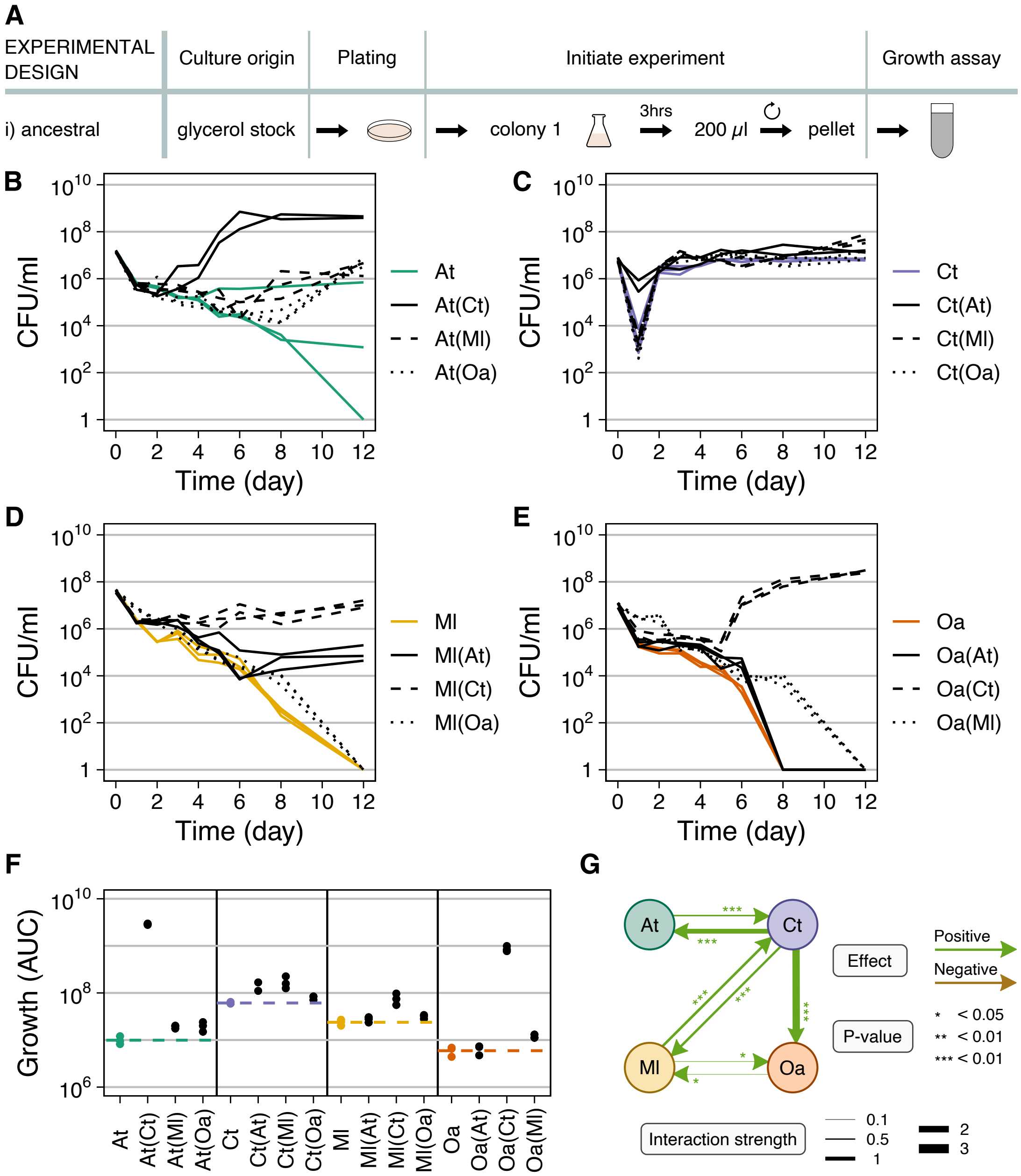


Figure S4: Comparison of ancestral mono- and pairwise co-cultures, adapted from[34]. (A) Glycerol stock of ancestral isolate was grown alone 3 hours to exponential phase, then washed and resuspended in MWF. (B=E) Population size quantified in colony-forming units per milliliter over time for mono-cultures (in color) and pairwise co-cultures (in black; co-culture partner indicated in brackets). In the cocultures, each species could be quantified separately by selective plating. Each panel shows the data for 1 species: (B) *A. tumefaciens* (At), (C) *C. testosteroni* (Ct), (D) *M. liquefaciens* (Ml) and (E) *O. anthropi* (Oa). (F) AUC in B=E. Dashed lines indicate the mean of the mono-cultures, shown in color.

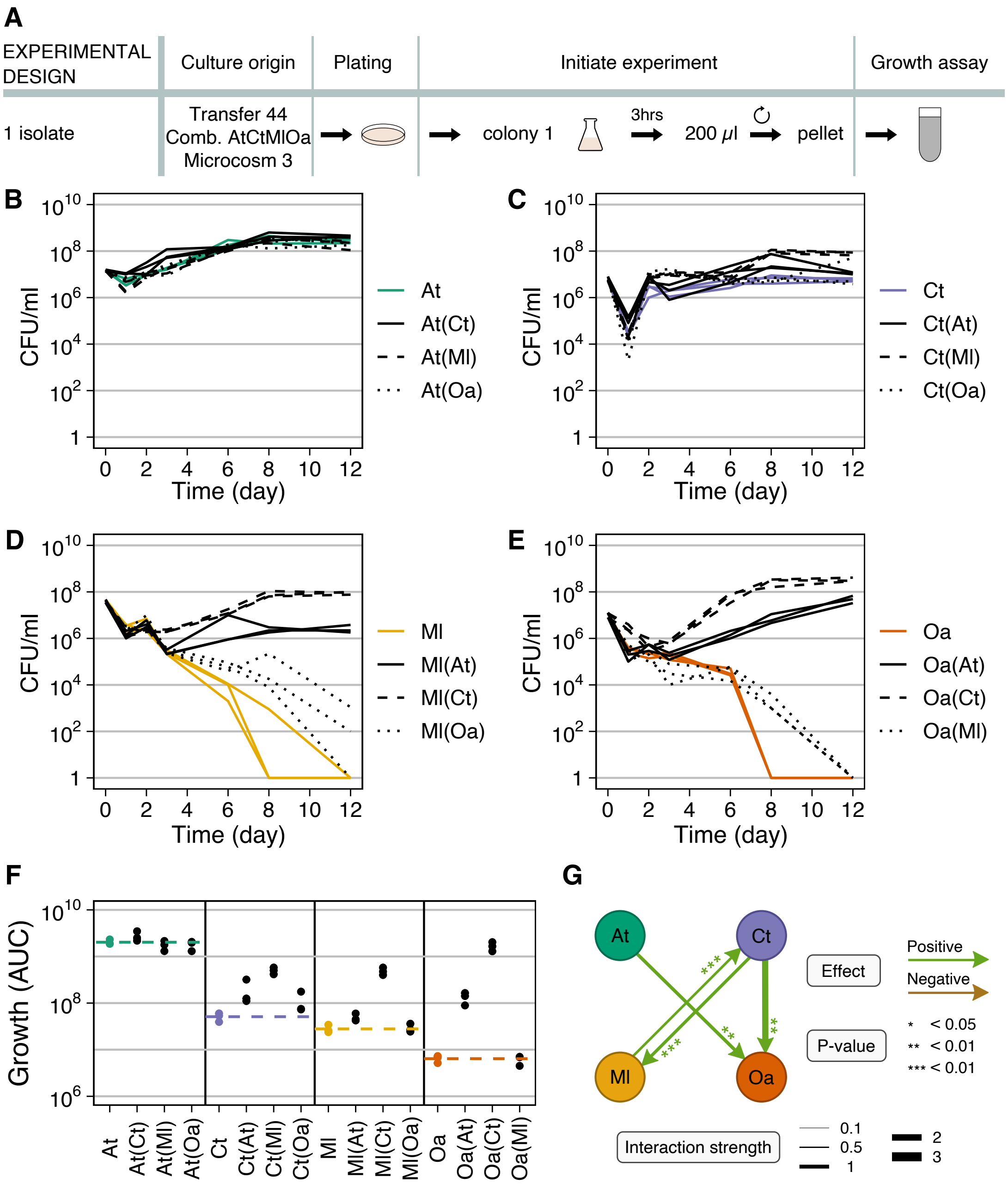


Figure S5: Comparison of co-evolved microcosm 3 mono- and pairwise co-cultures. (A) One evolved isolate of each species was randomly picked and grown alone 3 hours to exponential phase, then washed, resuspended and mixed in equal proportions in MWF. (B=E) Population size quantified in colony-forming units per milliliter over time for mono-cultures (in color) and pairwise co-cultures (in black; co-culture partner indicated in brackets). In the co-cultures, each species could be quantified separately by selective plating. Each panel shows the data for 1 species: (B) *A. tumefaciens* (At), (C) *C. testosteroni* (Ct), (D) *M. liquefaciens* (Ml) and (E) *O. anthropi* (Oa). (F) AUC in B=E. Dashed lines indicate the mean of the mono-cultures, shown in color. Statistical significance and interaction strengths data are shown in Dataset S2.

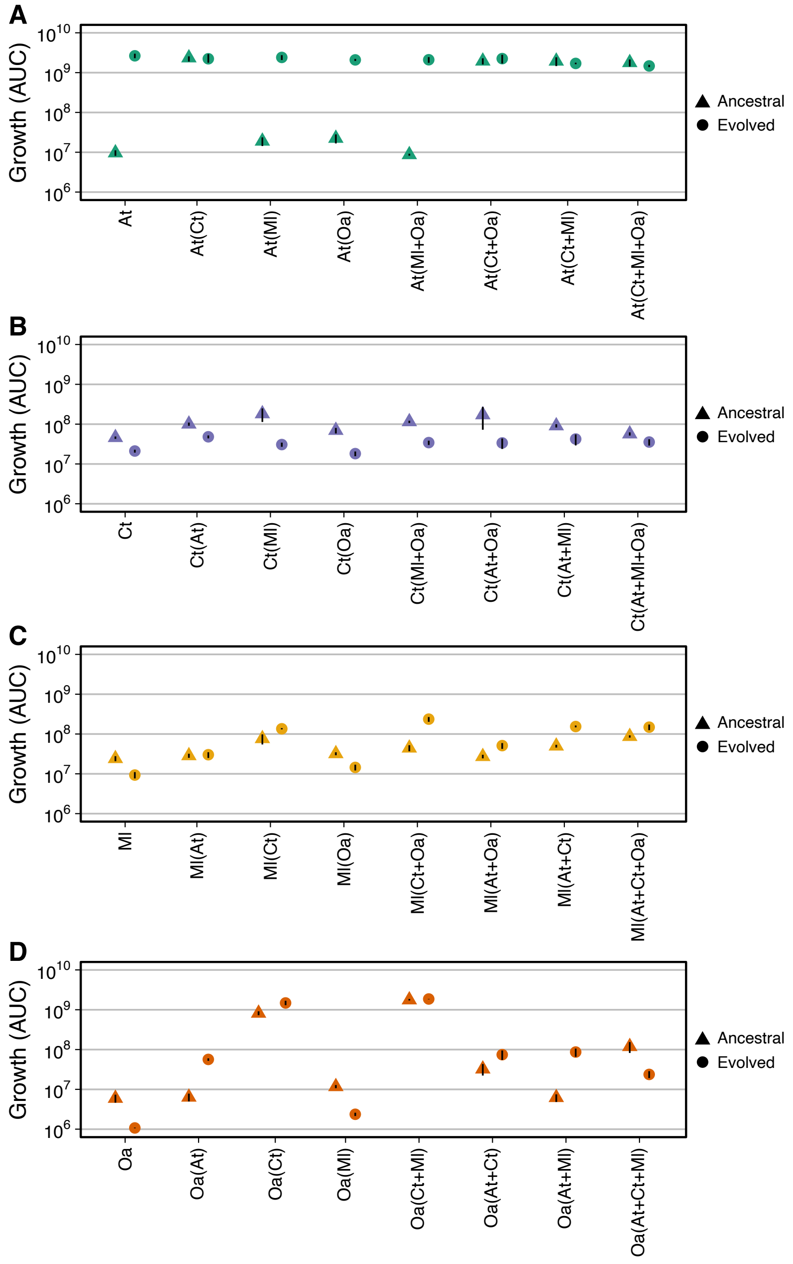


Figure S6: AUC comparison of ancestral species and those evolved in CAtCtMlOa, microcosm 3, including mono- and co-cultures treatments for (A) *A. tumefaciens*, (B) *C. testosteroni*, (C) *M. liquefaciens*, and (D) *O. anthropi*. Evoled strains were co-culured with isolates from the same microcosm and ancestral strains were co-cultured with other ancestors.

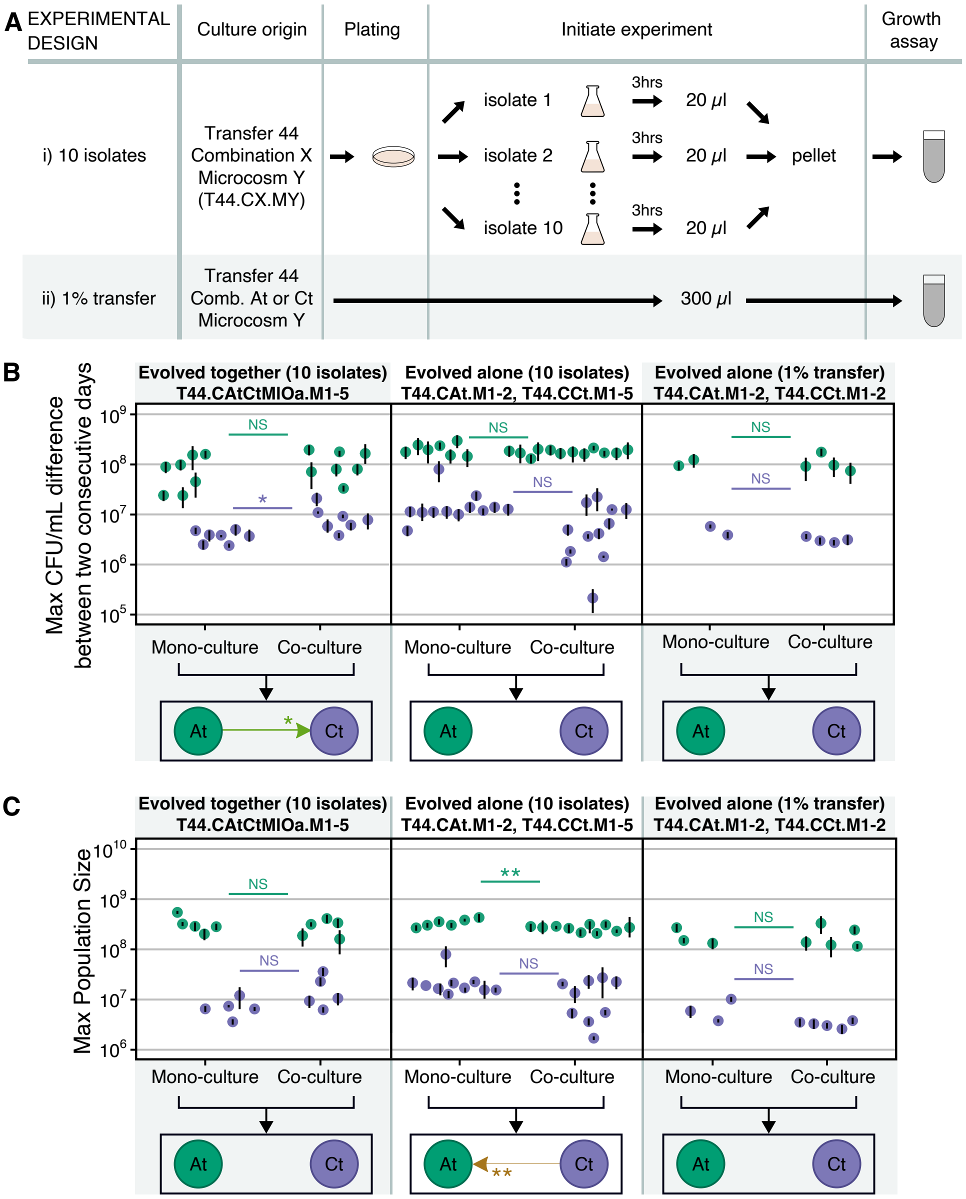


Figure S7: Interactions based on maximum growth rate and maximum population size. A) Protocols for growth assays, matching those in Fig. 3A. (B-C) Interactions between *A. tumefaciens* and *C. testosteroni* based on maximum growth rate quantified as the maximal CFU/ml difference between two consecutive days (B) or maximum population size (C), either evolved together (first column, CAtCtMlOa) or evolved alone (2nd and 3rd column, CAt and CCt, protocols i and ii from panel A) during 8-day growth assays. Other details are as in Fig. 3C.

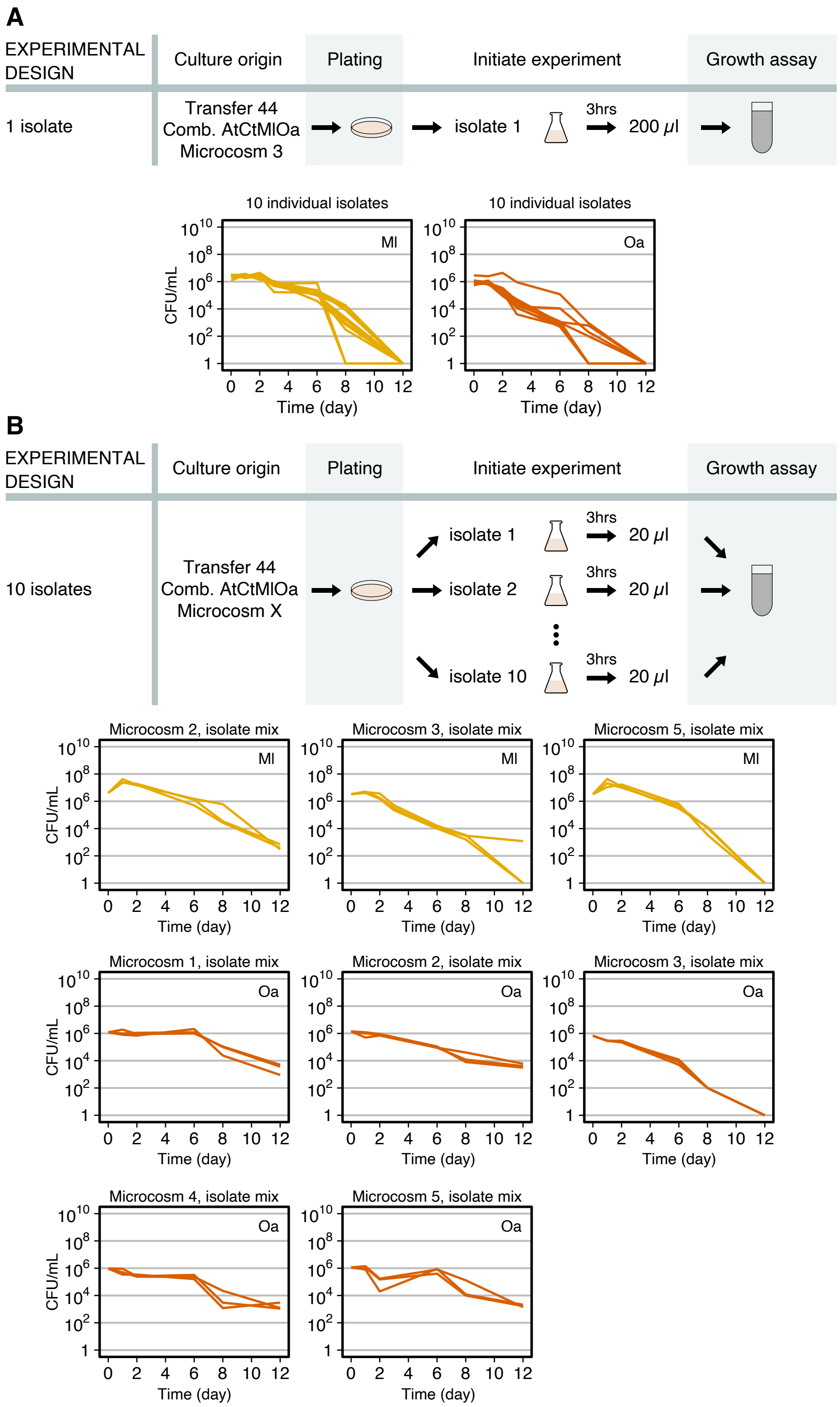


Figure S8: Mono-culture growth curves of evolved *M. liquefaciens* or *O anthropi* from transfer 44, CAtCtMlOa during 12-day growth assays. Conditions and microcosms are indicated above each graph. (A) One isolates was randomly picked and grown alone 3 hours to exponential phase, then washed and resuspended in MWF. Each growth curve represents one of 10 such isolates. (B) Ten evolved isolates were randomly picked and grown alone 3 hours to exponential phase, then washed, resuspended as a mixed culture in MWF. Each panel shows triplicates of isolates the same condition and microcosm.

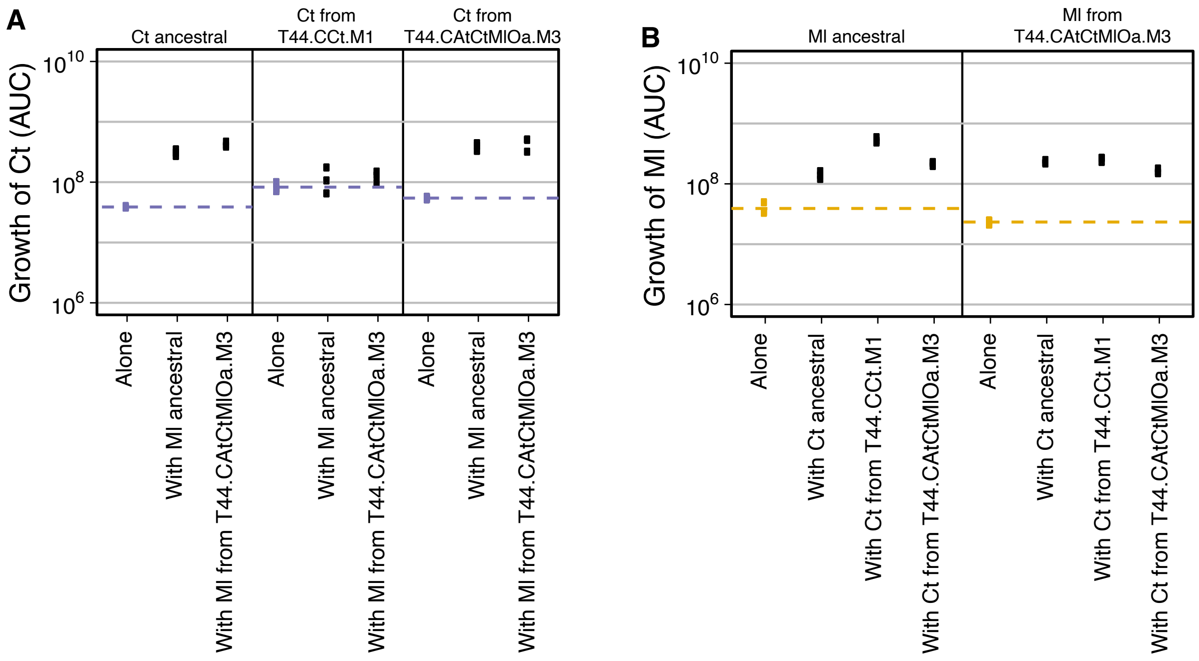


Figure S9: Interactions between *Ct* and *Ml*. (A) Growth of different *Ct* isolates (ancestral, evolved alone or evolved with the three others) alone or in co-culture with different *Ml* isolates (ancestral or evolved with the three others). (B) Growth of different *Ml* isolates alone or in co-culture with different *Ct* isolates. Community-evolved *Ct* and *Ml* were isolated from the same microcosm. Ancestral and community-evolved *Ct* and *Ml* all had positive effects on one another, but the positive effects did not increase between the isolates of the two species coming from the same microcosm, suggesting that at least in this microcosm, *Ct* and *Ml* did not evolve stronger mutualism.

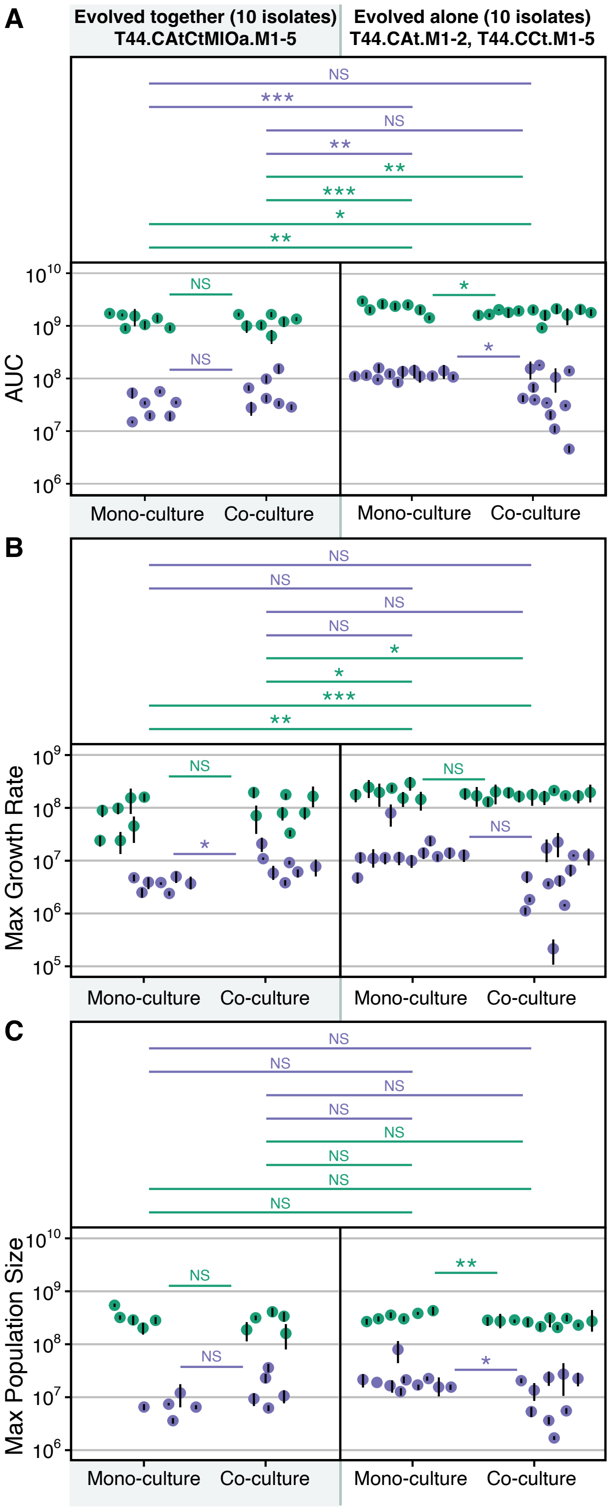


Figure S10: Inter-group comparison from Fig. 3C. The data show interactions between *A. tumefaciens* and *C. testosteroni* co-evolved (first column) or mono-evolved (second column) during 8-day growth assays. The first row measures the AUC of their growth curves during 8-day growth assays. The second and third row measure their maximum growth rates and maximum population size reached during these growth assays.

**A**

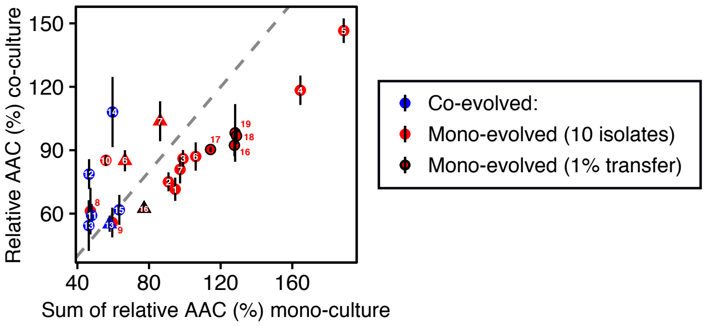


1a) We performed linear regression and calculated the estimated value of the coefficient as well as its standard error (these data are needed to calculate t-test):

- Linear regression, coefficient = 0.091, std error= 0.176, p-value < 0.05 *
- Linear regression, coefficient = 1.471, std error= 0.266, p-value < 0.001 ***
- Linear regression, coefficient = 0.091, std error= 0.157, p-value < 0.001 ***

1b) Next, we calculated p-value against the null hypothesis H0(slope) = 1:

- T-test, t = -5.170 , p-value < 0.05 *, following Bonferroni correction
- T-test, t = 1.771, p-value = 0.214, following Bonferroni correction
- T-test, t = 3.258, p-value = 0.094, following Bonferroni correction

2) We performed linear model to compare between groups:

- vs. Linear model with biological replicates as random factor, t = 2.418 , p-value = 0.056, following Bonferroni correction
- vs. Linear model with biological replicates as random factor, t = 6.450, p-value = 0.000236 ***, following Bonferroni correction
- vs. Linear model with biological replicates as random factor, t = 0.933, p-value = 0.732, following Bonferroni correction

**B**

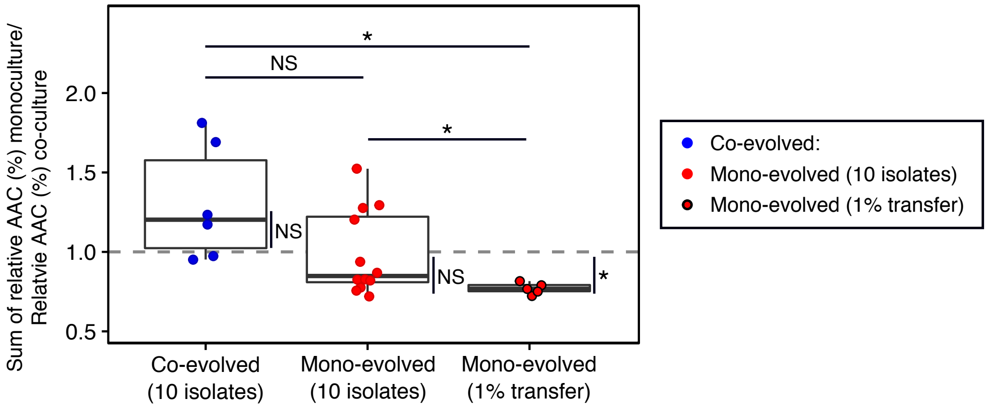


1) We calculated p-value against the null hypothesis H0(median) = 1:

- Sign-test, is the median greater than H0(median) = 1 ?, p-value = 0.3437
- Sign-test, is the median lower than H0(median) = 1 ?, p-value = 0.1938
- Sign-test, is the median lower than H0(median) = 1 ?, p-value = 0.0392 *

2) We performed t-test to compare between groups:

- vs. T-test, t = -1.9306 , p-value = 0.181, following Bonferroni correction
- vs. T-test, t = 3.5878, p-value = 0.03028 *, following Bonferroni correction
- vs. T-test, t = 2.6955, p-value < 0.0392 *, following Bonferroni correction

Figure S11: Results of statistical analysis of the additive null model to degradation efficiency in Fig. 5C). (A) Linear model. (B) T-test. Co-evolved is from species combination CAtCtMlOa and mono-evolved from CAt or CCt.

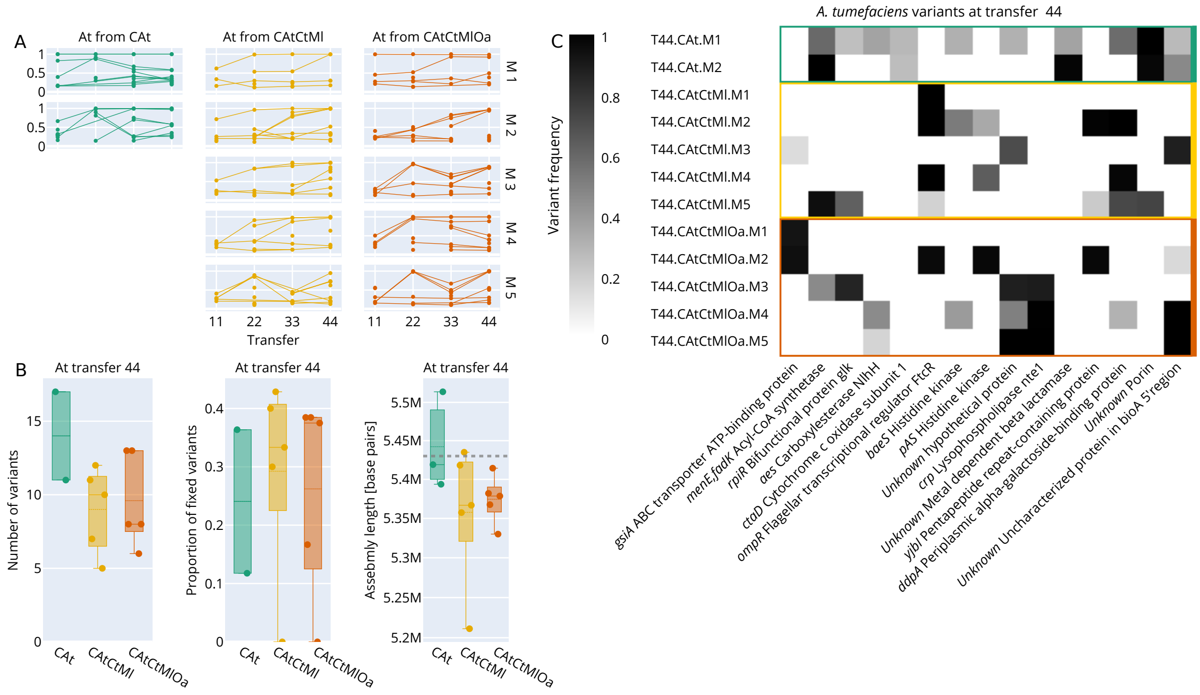

Figure S12: (A) Variant frequency trajectories in all *A. tumefaciens* populations. (B) Number of variants found in each *A. tumefaciens* population (left). De-novo long-read assembly lengths of selected isolates. Dashed line represents assembly length of the ancestor (middle). Proportion of variants that reached fixation (right). (C) Mutated genes with protein annotation that were found in at least two *A. tumefaciens* populations. The color indicates the frequency of the mutated allele.

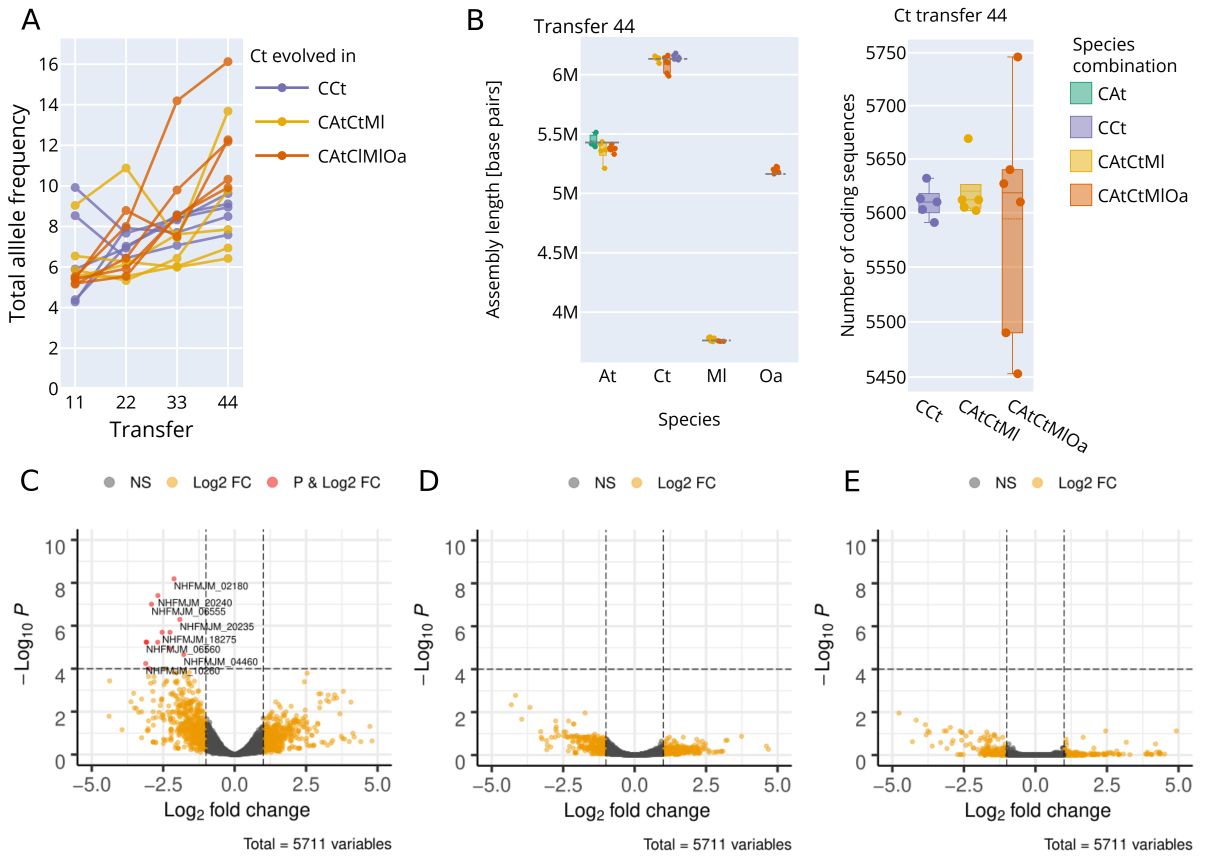

Figure S13: (A) Total allele frequency for *C. testosteroni*. (B) Long-read assembly lengths of isolates from transfer 44 with the dashed line representing the assembly length of the ancestor (left). Number of coding sequences per assembly (right). (C) Gene expression for *C. testosteroni* evolved under condtition 2 compared to ancestor. (D) Gene expression for *C. testosteroni* evolved under condtion 3 compared to ancestor. (E) Gene expression for *C. testosteroni* evolved under CAtCtMlOa compared to ancestor.

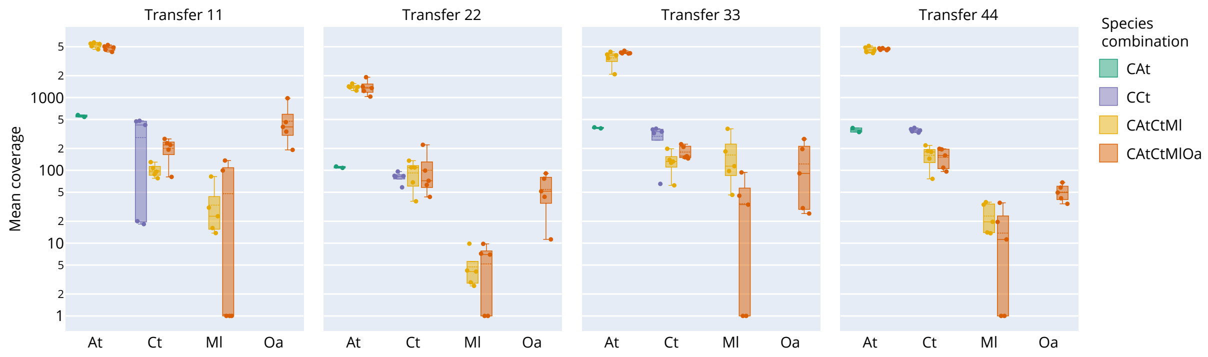

Figure S14: Mean Illumina coverage.

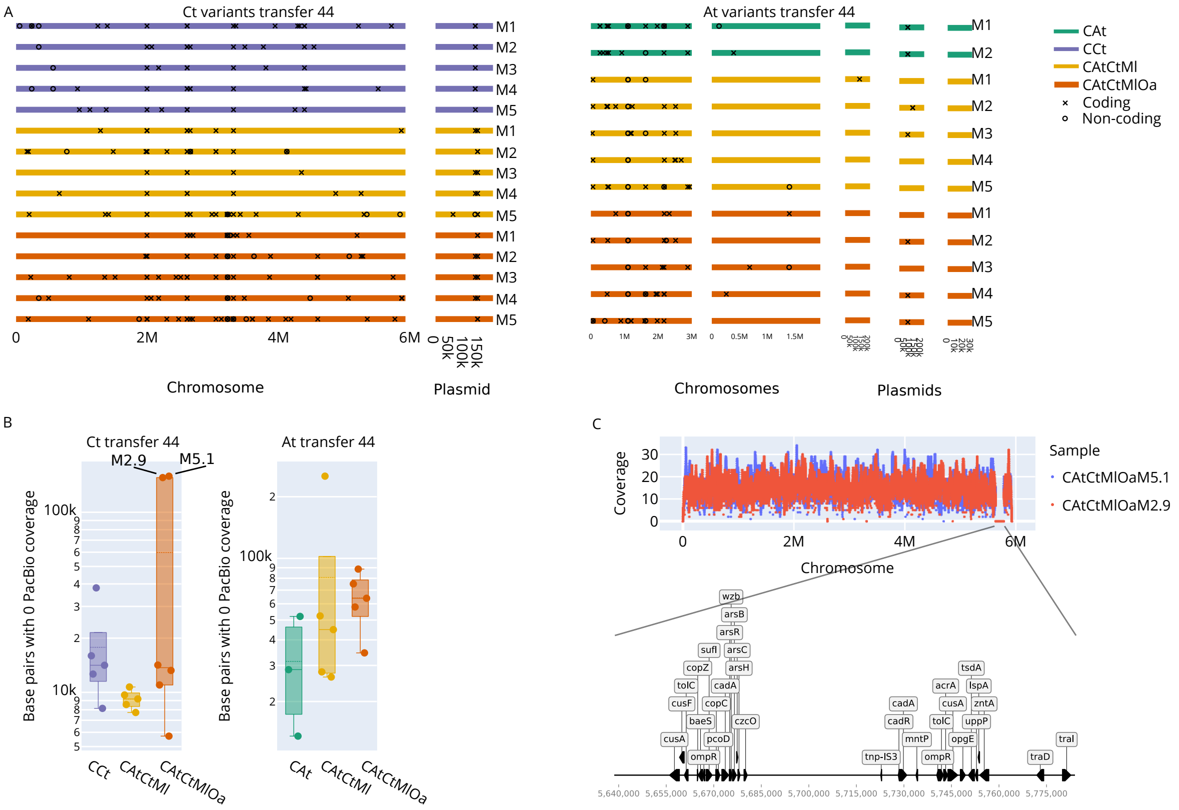


Figure S15: (A) Positions of variants across all frequencies identified from the Illumina data from the last transfer. (B) Base pairs with zero-coverage when aligning corrected PacBio reads to the reference genome. (C) PacBio coverage for two Ct isolates of CAtCtMlOa showing large deletion (top). Annotation of deleted sequence (bottom).

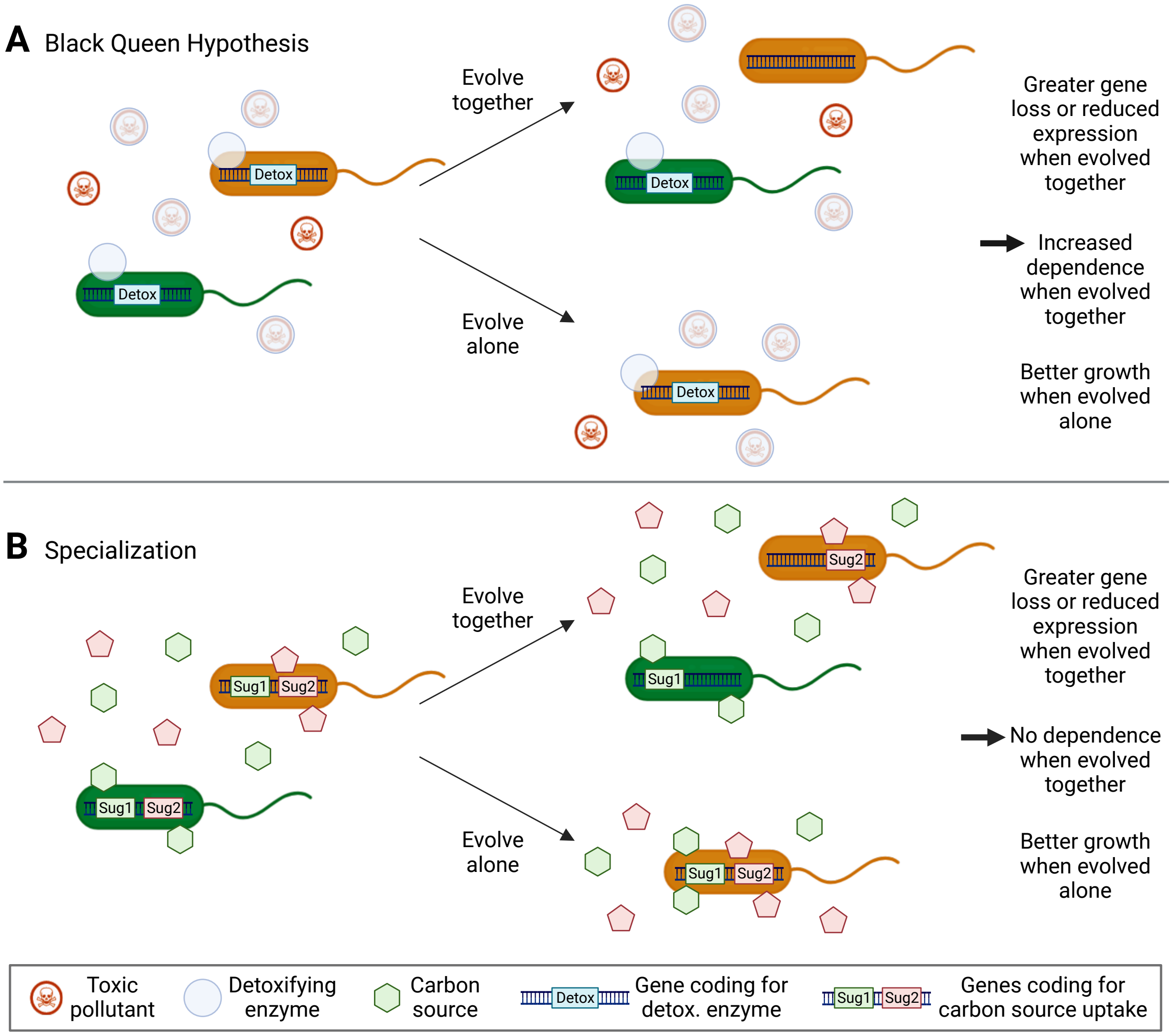


Figure S16: BQH and specialization make similar predictions. (A) The BQH predicts that species evolved together in community should lose traits coding for public goods, like detoxification genes either by deletions or mutations leading to reduced gene expression. Such losses should not be observed when evolving alone. Species evolved together should therefore grow significantly worse alone and depend on the partner species for survival. (B) The evolution of specialization predicts similar trait loss when evolving together and should similarly grow best when evolved alone, but species evolved in community should not depend on their partners to grow alone (black arrows on the right). Initially both species can take up both carbon sources but with a preference for one or the other. After evolution alone, the orange species can take up both efficiently. Generated using Biorender.

| Gene ID | Gene | Product | Log2 fold change | -Log10P |
|---|---|---|---|---|
| NHFMJM_02180 | Unknown | hypothetical protein | -2.122160 | 8.186240 |
| NHFMJM_04460 | Unknown | hypothetical protein | -1.783635 | 4.657574 |
| NHFMJM_06555 | Unknown | Putative lipoprotein | -2.909484 | 7.003442 |
| NHFMJM_06560 | *rpoE* | RNA polymerase, sigma-24 subunit, ECF subfamily | -3.096592 | 5.238158 |
| NHFMJM_10260 | *raiA* | Ribosomal subunit interface protein | -3.110503 | 4.240304 |
| NHFMJM_18275 | Unknown | 3-demethylubiquinone-9 3-methyltransferase | -2.537547 | 5.694750 |
| NHFMJM_20235 | Unknown | DUF937 domain-containing protein | -1.919700 | 6.292688 |
| NHFMJM_20240 | Unknown | Inosine-5'-monophosphate dehydrogenase | -2.685720 | 7.404975 |

Table S2: Differentially expressed genes in *C. testosteroni* evolved alone (CCt).
